# HSP90 buffers deleterious genetic variations in *BRCA1*

**DOI:** 10.1101/2024.11.15.623783

**Authors:** Brant Gracia, Patricia Montes, Min Huang, Junjie Chen, Georgios Ioannis Karras

## Abstract

Protein-folding chaperone HSP90 buffers genetic variation in diverse organisms, but the clinical significance of HSP90 buffering in disease remains unclear. Here, we show that HSP90 buffers mutations in the BRCT domain of BRCA1. HSP90-buffered *BRCA1* mutations encode protein variants that retain interactions with partner proteins and rely on HSP90 for protein stability and function in cell survival. Moreover, HSP90-buffered BRCA1 variants confer PARP inhibitor resistance in cancer cell lines. Low-level HSP90 inhibition alleviates this resistance, revealing a cryptic and mutant-specific HSP90-contingent synthetic lethality. Hence, by stabilizing metastable variants across the entirety of the BRCT domain, HSP90 reduces the clinical severity of BRCA1 mutations allowing them to accumulate in populations. We estimate that HSP90 buffers 11% to 28% of known human BRCA1- BRCT missense mutations. Our work extends the clinical significance of HSP90 buffering to a prevalent class of variations in *BRCA1*, pioneering its importance in cancer predisposition and therapy resistance.

## INTRODUCTION

Despite the advent of high-throughput functional assays for evaluating mutation pathogenicity,^1–6^ predicting the clinical course of genetic diseases in humans remains difficult.^7^ A significant unaddressed issue is the conditionality of genetic variation whereby genetic modifiers, environmental factors, or lifestyle choices alter the clinical presentation of disease.^8–10^ Indeed, incomplete mutation penetrance and variable clinical severity (expressivity) pervade a host of human genetic disorders.^11^ Hence, defining mechanisms that broadly modify the biological effects of genetic variation is expected to enhance our understanding of diverse diseases and aid in developing strategies to predict and prevent pathology.^12–21^

Insights into the conditionality of genetic variation have emerged from research in the field of protein folding. Proteins are polypeptide chains that typically fold into well-defined three-dimensional structures to function. Because a protein’s native structure is only marginally more stable than non-native conformations,^22–24^ small variations in solution or temperature conditions can profoundly impact protein stability.^25–27^ Furthermore, changes in amino acid sequence induced by mutations can destabilize proteins in the face of such environmental fluctuations. Functional assays in cells suggest that ∼30-60% of all pathogenic missense mutations destabilize protein structure.^2,28–30^ Moreover, the percentage of destabilizing pathogenic variants can reach values upwards of 90% depending on the gene or gene region.^31^ Still, many proteins can tolerate genetic variation,^32^ suggesting cells utilize specific buffering (i.e., salvaging) mechanisms to support the activity of mutant proteins.^33^ Identifying buffering mechanisms that can explain the conditional effect of many clinically-relevant mutations would significantly advance our ability to predict the course of complex diseases.

The protein-folding chaperone heat shock protein 90 (HSP90) is a quintessential buffering mechanism in biology. HSP90 is essential in most living organisms, comprising a chaperone reservoir greater than the requirements for maintaining cell viability and survival.^34,35^ Excess HSP90 provides a protein folding buffer that safeguards proteostasis in the face of proteotoxic stress and can influence the course of pathologies driven by the accumulation of misfolded proteins.^8,36–38^ Moreover, excess HSP90 buffers genetic variation across diverse organisms, thereby potentially altering the course of evolutionary processes.^39–44^ Importantly, seemingly benign proteotoxic stressors tax HSP90 buffering activity within the cell. As a result, proteins that rely on HSP90 can lose function. This profound environmental sensitivity allows HSP90 to mediate gene-environment interactions.^39–44^ A recent investigation probing the role of HSP90 in yeast domestication underscores the importance of HSP90 in evolution as mediator of ecologically relevant gene-by-environment interactions.^45^ Moreover, HSP90 has been shown to buffer mutations in humans by stabilizing the encoded proteins, as exemplified by a case of twins with Fanconi Anemia.^46^ However, the prevalence of HSP90-buffered variants in populations, their importance for cancer predisposition, impact on response to therapy, and their overall significance for human health remain to be determined.

Here, we expand the significance and define the mechanism of HSP90 buffering in relation to pathogenic variations in the *BRCA1* tumor suppressor, the poster child gene of precision medicine.^47–49^ HSP90-buffered variants of the BRCT domain in BRCA1 (BRCA1-BRCT) exhibit molecular properties characteristic of partially destabilized proteins and rely on HSP90 for function. HSP90- buffered BRCA1 variants retain interactions with partner proteins and support cell survival. HSP90 offsets the destabilizing effects of mutations by protecting the encoded BRCA1 variants from degradation. In cancer cells, HSP90 buffering promotes resistance to the PARP inhibitor olaparib similar to wild-type cells. Low-level, non-toxic concentrations of HSP90 inhibitors synergize with PARP inhibition to kill cancer cells expressing HSP90-buffered *BRCA1* mutations but not wild-type cells, highlighting a novel BRCA1 polytherapy approach based on targeting HSP90’s buffering function. HSP90-buffered BRCA1-BRCT variants account for ∼11-28% of missense variants we tested across diverse datasets and confer delayed breast cancer onset in humans. Our results delineate the criteria for HSP90 buffering in *BRCA1* and warrant their use to stratify cancer patients with HSP90-buffered *BRCA1* mutations for diagnosis, prognosis, and tailored management, including combination therapies with HSP90 inhibitors.

## RESULTS

### Essential BRCA1 function evaluated using degron mediated protein depletion

Mutations in tumor suppressor *BRCA1* greatly increase cancer predisposition in human carriers because the encoded protein is critical for the homology-directed repair of DNA double strand breaks.^3,47,50^ Because this function is essential in vertebrates, *BRCA1*-null cells exhibit dramatically decreased viability and CRISPR/Cas9 knock-in efficiency,^51,52^ precluding the study of pathogenic variations at the endogenous *BRCA1* locus. Hence, we employed the degradation tag (dTAG) system, which leverages cell-permeable heterobifunctional degraders to conditionally deplete the endogenous BRCA1 protein in mammalian cells.^53,54^

To evaluate the effect of BRCA1 degradation, we tested an oral squamous cell carcinoma cell line because previous reports suggest BRCA1 expression correlates with poor prognosis (Cal 27)^55,56^ as well as a non-cancer human embryonic kidney 293-derived cell line (HEK293A). We used CRISPR/Cas9 to knock-in FKBP12^F36V^ in frame with the endogenous *BRCA1* locus, generating a chimeric protein that can be conditionally degraded upon addition of dTAG compound^53,54^ (Figures 1A and 1B). We validated the derived homozygous knock-ins by PCR and log-read sequencing of the engineered *BRCA1* locus (Figures S1A, S1B, and Methods). Two independent dTAG-BRCA1 clones (C2 and C3) exhibited no significant difference in growth rate as compared to the parental cell line (Figures S1C), despite expressing the chimeric FKBP12^F36V^-2xHA-BRCA1 protein—referred to as dTAG-BRCA1 henceforth (Figures S1D). Thus, our approach generated homozygous knock-in cell lines expressing chimeric dTAG-BRCA1 protein that fully supports the essential function of BRCA1 (BRCA1^dTAG/dTAG^).

**Figure 1.**
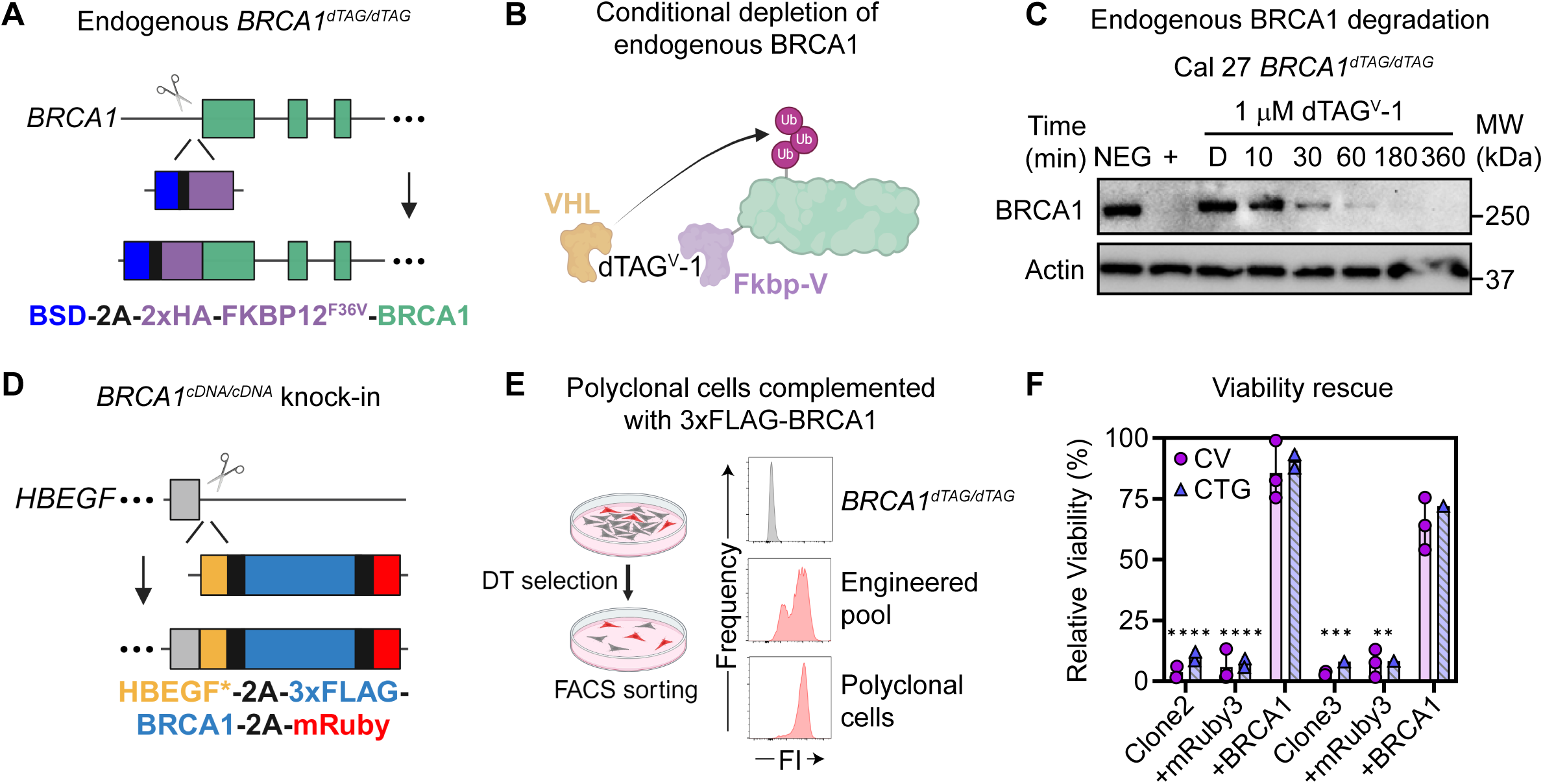
Essential BRCA1 function evaluated using degron mediated protein depletion. (A, B) Genome engineering of the endogenous *BRCA1* gene locus to express FKBP12^F36V^-BRCA1 chimera in the homozygous state (*BRCA1^dTAG/dTAG^*). A Blasticidin resistance gene (BSD) is expressed upstream of BRCA1 separated by a 2A ribosome skipping sequence to facilitate knock-in selection. Shown in (B) is recruitment of the von Hippel-Lindau (VHL) E3 ligase complex (shown as one blob for simplicity) upon addition of the heterobifunctional dTAG^V^-1 (dTAG herein) molecule. Ub, ubiquitin. HA, hemagglutinin tag. Fkbp-V, FKBP12^F36V^. (C) Conditional depletion of dTAG-BRCA1 chimera in Cal 27 cells after treatment with 1 µM dTAG as measured by Western blotting. Negative control (indicated by “NEG“) was treated with dTAG-NEG compound which cannot promote degradation, and positive control (indicated by “+“) was treated with dTAG overnight. MW, molecular weight. kDa, kiloDaltons. D, vehicle (DMSO). (D, E) Genome engineering enhanced by diphtheria toxin (DT) to complement loss of dTAG-BRCA1 using BRCA1 cDNA transgene. HBEGF* contains an exon 4 mutant that confers DT resistance and exon 5. BRCA1 and a fluorescent reporter are interspaced with 2A ribosome skipping sequences. Example flow cytometry results are shown at each stage of engineering for knock-in of 3xF-BRCA1- 2A-mRuby3. 3xF, triple FLAG tag. FACS, Fluorescence-activated cell sorting. FI, fluorescence intensity. (F) Cell survival assays to quantify *BRCA1*-dependent viability in two Cal 27 *BRCA1^dTAG/dTAG^* clones engineered with mRuby3 or 3xF-BRCA1-2A-mRuby3. Relative viability (%) calculated by normalizing 1 µM dTAG treated cells to dTAG-NEG control treatments for each cell line, separately. Quantification of cell survival using crystal violet staining (CV, purple circles) mimicked cell viability effects as measured by CellTiter-Glo® (CTG, blue triangles). Statistical significance was determined using Kruskal-Wallis ANOVA test (F). \*\*\*\**p*≤0.0001. \*\*\**p*≤0.001. \*\**p*≤0.01. Data are presented as mean ± standard deviation values from at least two independent experiments or two independent clones.

Next, we tested the stability of dTAG-BRCA1 upon treatment with degradation-inducing compound dTAG^V^-1 (dTAG herein). In one HEK293A clone and five Cal 27 clones, dTAG treatment resulted in BRCA1 protein degradation but did not affect the levels of BRCA1 expressed in the parental (P+) non-engineered cells (Figure S1E). In addition, dTAG-induced BRCA1 depletion was rapid (half-life of ∼15 minutes) in both HEK293A and Cal 27 cells (Figures 1C and S1F). In contrast, dTAG-BRCA1 expression was unaffected by a stereochemical isomer unable to activate the degron (dTAG^V^-1-NEG, dTAG-NEG herein), highlighting the specificity of BRCA1 depletion using dTAG. Furthermore, dTAG- BRCA1 depletion was irreversible over six days of dTAG treatment (Figure S1G). These results demonstrate the rapid, robust, and conditional depletion of endogenous dTAG-BRCA1 protein.

Next, we tested if loss of BRCA1 inhibits the survival of *BRCA1^dTAG/dTAG^* cells, as is expected from *BRCA1* inactivation in other systems.^3,51^ In HEK239A and Cal 27 cells, we observed a dose-dependent reduction in cell proliferation with increasing concentrations of dTAG (Figures S1H and S1I). In addition, BRCA1 depletion reduced cell proliferation in all homozygous dTAG-BRCA1 clones but not in a heterozygous clone that retained the wild-type allele (Figure S1J, C4*). These results demonstrate that BRCA1 depletion induces a substantial growth defect in *BRCA1^dTAG/dTAG^* cells.

We next developed a complementation approach to stably express full-length *BRCA1* transgenes in *BRCA1^dTAG/dTAG^* cells. Our approach was based on a previous Cas9-based strategy to integrate transgenes into the *HBEGF* safe-harbor locus using negative selection by treatment with Diphtheria toxin (DT).^57^ We engineered donor templates to integrate an *HBEGF* variant that confers resistance to DT (HBEGF*) while simultaneous introducing 3x-FLAG tagged full-length BRCA1 (3xF-BRCA1 herein) cDNA followed by a 2A ribosome skipping sequence and mRuby3 fluorescent protein (Figure 1D). This strategy enables DT selection and FACS sorting of polyclonal pools of BRCA1 positive fluorescent cells (Figure 1E). Fluorescence expression was stable for up to 38 passages indicating the polyclonally engineered cells retained long-term stability in BRCA1 expression (Figure S1K). Integration of the wild-type *BRCA1* transgene completely restored clonogenic survival (crystal violet staining) and viability (CellTiter-Glo® assay) in both Cal 27 and HEK293A backgrounds as compared to integration of the fluorophore alone (Figures 1F, S1L, and S1M). Altogether, these results demonstrate a robust complementation system to evaluate the function of candidate HSP90-buffered and non-buffered BRCA1 variants.

### HSP90-engaged BRCA1-BRCT variants retain protein partner binding and support cell survival

The BRCT domain of BRCA1 (BRCA1-BRCT) scaffolds phosphorylated partner proteins (pSXXF) during DNA repair. Pathogenic BRCA1-BRCT variations exhibit dramatic variability in binding to protein-folding chaperone HSP90 and a related co-chaperone HSP70.^31^ Strong HSP70 binding to BRCA1-BRCT variants correlated with severe disruption to domain folding and function, consistent with HSP70’s role in binding unfolded hydrophobic polypeptides. Hence, HSP90-engaged variants were classified as moderately HSP90-bound variants that did not strongly bind to HSP70. By comparing HSP90 binding magnitude with available functional data,^3,5,58^ we observed that moderately HSP90-engaged BRCA1-BRCT variants tend to retain BRCA1 activity compared to other variant groups (Figure 2A). Considering the importance of protein partner binding to the BRCA1-BRCT domain in tumor suppression,^59^ we speculated that moderate HSP90 binding reflects partially folded variants that retain PPIs similar to the wild-type.

**Figure 2.**
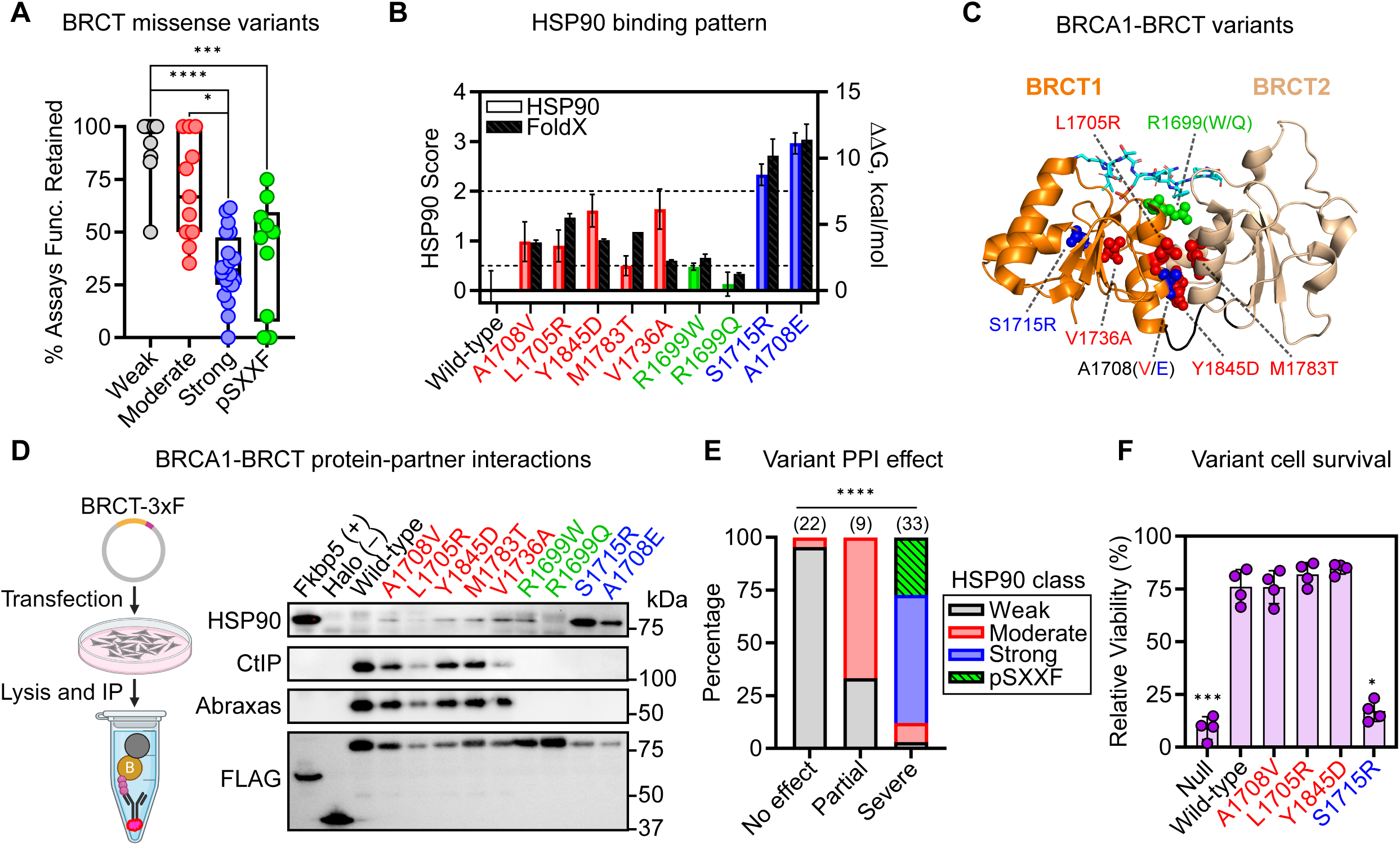
HSP90-engaged BRCA1-BRCT variants retain protein partner binding and support cell survival. (A) Moderate HSP90 binding to BRCA1-BRCT variants correlates with maintenance of protein function. Shown is the percent of functional assays for each variant that reported “functional” phenotypes. Variants are binned by the magnitude of HSP90 binding or phosphopeptide (pSXXF) coordinating residues.^58^ Only variants with three or more assays reported are shown. Func., function. (B – C) HSP90 binding magnitude of representative variants from each class (left y-axis) and the predicted destabilizing effect of mutation as measured by FoldX^113^ (right y-axis). Panel (C) shows the physical location of each mutated residue in the BRCT crystal structure^114^ (pSXXF binding peptide from the structure is shown in cyan). HSP90 moderate, red; pSXXF, green; HSP90 strong, blue. (D) CoIP approach to detect BRCA1-BRCT protein-partner interactions in HEK293T cells using truncation construct 1,314-BRCT C-terminally tagged with 3x-FLAG (BRCT-3xF). Schematic at the left shows pull-down approach using agarose-linked anti-FLAG antibodies. Purple spheres represent 3xFLAG epitope. Fkbp5 and HALO-tag proteins serve as HSP90 positive (+) and negative (–) controls, respectively. kDa, kiloDaltons. (E) Moderate HSP90 binding to BRCA1-BRCT variants correlates with maintenance of PPIs. Each bin shows the severity effect of mutation on BRCT pSXXF binding (no effect, partial, severe).^61^ Variants were assigned an HSP90 binding class or pSXXF. The number of variants in each bin is shown in parenthesis above the stacked bar. (F) BRCA1 mutant cell survival phenotypes. The relative viability (%) was quantified by normalizing to dTAG-NEG treated cells for each cell line separately. Statistical significance was determined using Kruskal-Wallis ANOVA test (A and F) or Fisher’s exact test (E). \*\*\*\**p*≤0.0001. \*\*\**p*≤0.001. \*\**p*≤0.01. \**p*≤0.05. Data are presented as mean ± standard deviation values from at least two independent experiments.

To test this hypothesis, we investigated a collection of BRCA1-BRCT variants^31^ several of which were engaged by HSP90 (red, Figures 2B and 2C). We specifically chose variants distal from the pSXXF binding site, including some previously described as hypomorphic.^31^ As controls, we included strongly destabilizing pathogenic variants that bind strongly to HSP90 and HSP70 (blue)^60–63^ and known pSXXF-disrupting variants (green).^60–62,64,65^ FoldX predictions on the destabilizing effect of mutations (ΔG_Mut_ – ΔG_WT_) qualitatively agreed with the magnitude of HSP90 binding indicating that weaker HSP90 binding reflects weaker effects on folding as previously described^31^ (Figure 2B). Indeed, *in vitro* purified proteins that bind weakly to HSP90 retained pSXXF binding (Figure S2A).^62^ Moreover, four moderately HSP90-engaged variants also retained pSXXF binding. In contrast, nearly all strong chaperone-bound variants exhibited significantly perturbed pSXXF binding, consistent with their strong destabilizing effects. These data suggest that HSP90-engaged BRCA1-BRCT variants typically retain the ability to interact with partner proteins *in vitro*.

To test for maintenance of protein-partner interactions in cells, we pulled-down transiently expressed C-terminally FLAG-tagged BRCT proteins from HEK293T cell lysates and detected the co-immunoprecipitated endogenous partner proteins using Western blotting (Figure 2D). As expected, BRCA1 partners CtIP and Abraxas co-immunoprecipitated with the wild-type BRCT.^66–68^ The HSP90 co-chaperone Fkbp5 served as a negative control for Abraxas and CtIP binding and a positive control for HSP90 binding. Strikingly, five out of five moderately HSP90-engaged variants retained binding to Abraxas and CtIP similar to the wild-type BRCA1 protein,^59^ in agreement with our hypothesis (Figure 2D). In contrast, variants S1715R and A1708E that interacted strongly with HSP90 did not bind CtIP and Abraxas, as expected for severely destabilized proteins. These results were reproducible in the contexts of an N-terminally tagged BRCT construct and the full-length BRCA1 protein (Figure S2B). Moreover, protein-partner interaction data^69^ revealed that variants moderately-engaged by HSP90 frequently retain partial BRCA1-BRCT PPIs (Figure 2E). Hence, moderately HSP90-bound BRCA1 variants often retain partner interactions in cells.

To test if HSP90-engaged BRCA1 variants also support cell viability, we stably expressed full-length 3xF-BRCA1 cDNA variants from the *HBEGF* locus of *BRCA1^dTAG/dTAG^* cells to measure changes in mutant-specific effects on cell survival. Upon dTAG-induced depletion of endogenous BRCA1, the variants were expressed at levels similar levels to the wild-type BRCA1 protein and dTAG treatment did not affect this expression (Figures S2C). Three moderately HSP90-bound BRCA1 variants supported cell survival similar to the wild-type allele in Cal 27 cells (Figure 2F). In contrast, the destabilized variant S1715R did not support viability, similar to the null. Furthermore, each cell population retained over 90% fluorescence over many generations, indicating that the variant cell populations are pure for the mutant of interest (Figure S2D). In addition, A1708V supported cell viability in two independent clones as measured by CellTiter-Glo® (Figure S2E). We demonstrated that our survival assay using crystal violet measured cell proliferation within a linear regime as long as cells were terminated before reaching full confluency (Figure S2F). Moreover, our survival assay was reproduced by CellTiter-Glo® viability assay (Figure S2G, R^2^=0.9, *p*<0.0001). We validated the BRCA1 complementation with each variant using PCR and Sanger sequencing (Figures S2H and S2I). Furthermore, we observed qualitatively similar effects of the BRCA1 variants on survival in HEK293A cells (Figures S2J and S2K). The maintenance of cell viability for A1708V and L1705R mutant was observed under different cell seeding and dTAG treatment conditions (Figure S2L). These results demonstrate that HSP90-engaged BRCA1 variants confer milder growth phenotypes, suggesting a direct role for HSP90 in buffering their deleterious effects.

### Pharmacological inhibition destabilizes HSP90-buffered BRCA1 variants

To test if HSP90 buffers mutations in BRCA1, we treated cells expressing different BRCA1 variants with HSP90 inhibitors and measured the effect on cell survival. Two HSP90 inhibitors were tested comprising unique chemical structures. NVP-HSP990 is a purine-containing compound in Phase I clinical trials while STA-9090 (Ganetespib) is a Resorcinol-containing compound currently in Phase III clinical trials.^70^ Both compounds bind the N-terminal domain of HSP90, blocking ATP binding and hydrolysis. We utilized these compounds to test if HSP90 is required for the essential function of HSP90-engaged BRCA1 variants.

Because HSP90 inhibitors are known to be toxic to cancer cells at high concentrations, we first identified non-toxic doses using cell survival in the context of wild-type BRCA1 by omitting dTAG treatment (Figure S3A). With these data, we treated Cal 27 mutant cells with a non-toxic dose of HSP90 inhibitor and measured the effect on cell survival. Strikingly, we observed mutant specific effects; cells dependent on BRCA1 mutants moderately-engaged by HSP90 (A1708V, L1705R, and Y1845D) were substantially more sensitive to HSP90 inhibition. Indeed, A1708V mutant cells decreased survival by 27% while wild-type cells were not significantly affected (Figure 3A). Moreover, similar mutant-specific effects in cell survival were observed when using HSP90 inhibitor STA-9090 in both Cal 27 and HEK293A cells, precluding cell-line specific effects of HSP90 stabilization (Figures S3B and S3C). These results suggest that HSP90 binding can suppress the growth defect of specific BRCA1 mutations.

**Figure 3.**
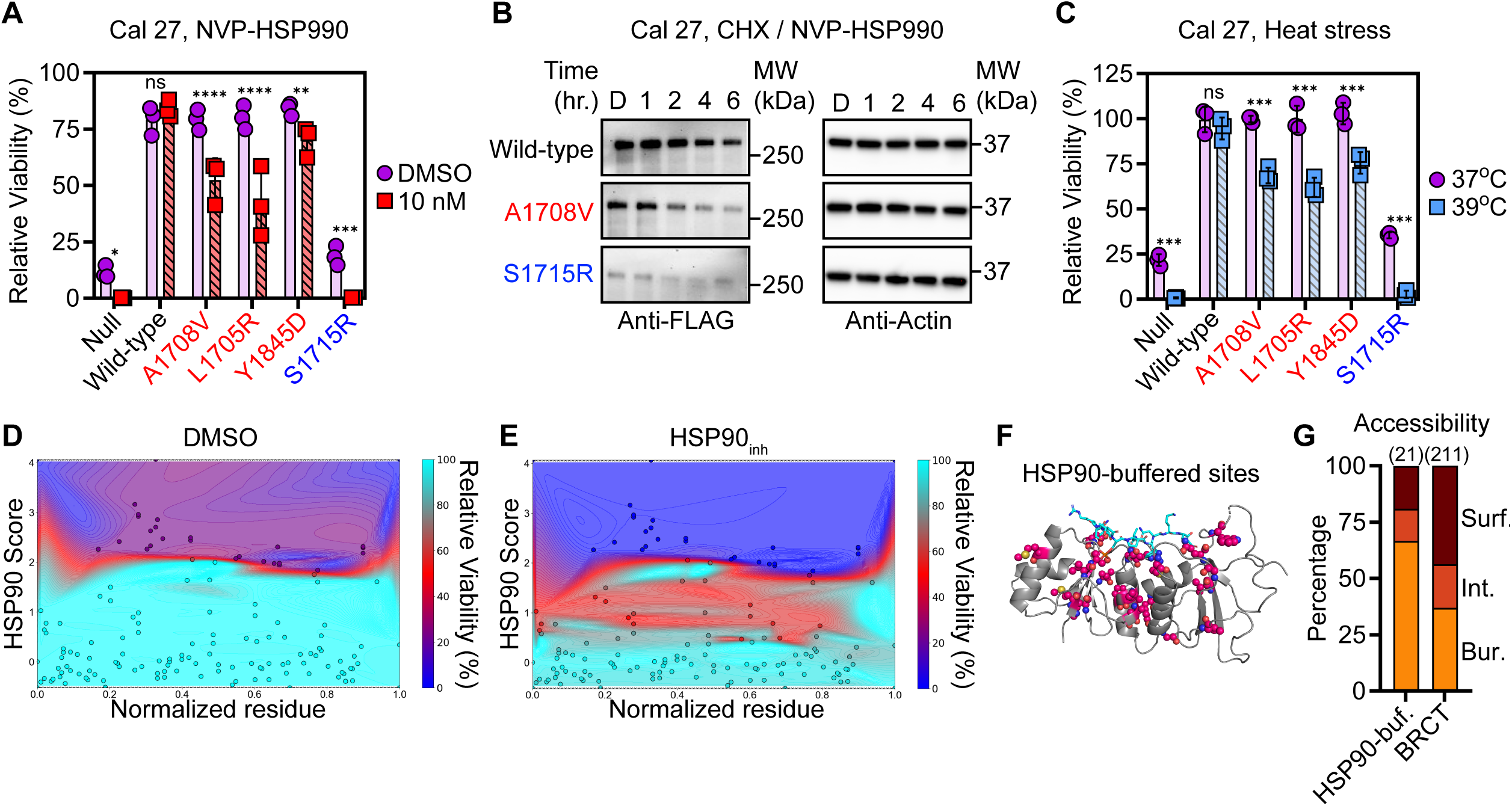
Pharmacological inhibition destabilizes HSP90-buffered BRCA1 variants. (A) HSP90-dependent survival of Cal 27 mutant cells expressing moderately (red) or strongly (blue) HSP90-engaged BRCA1 variants. Cells were treated with vehicle DMSO (purple circles) or with 10 nM NVP-HSP990 (red squares). Null cells were complemented with mRuby3 fluorophore alone. Relative viability (%) quantified by normalizing to dTAG-NEG treated cells for each cell line separately. (B) BRCA1 variant stability in the presence of HSP90 inhibitor. Cells were treated with 100 µg/mL CHX and 30 nM NVP-HSP990 for indicated number of hours followed by lysis and Western blotting to measure protein turnover. D, vehicle (DMSO). (C) Temperature sensitive Cal 27 mutant cells grown under basal conditions at 37 °C (purple circles) or heat stress at 39 °C (light blue squares). (D – E) Contour maps illustrating BRCA1-BRCT variant position, HSP90 binding score, and effect on survival under basal conditions (DMSO) and when HSP90 is inhibited (HSP90_inh_). Each data point (circles) contains three parameters. The x-axis shows BRCT residue normalized from 0 to 1 (1,650– 1,858). Y-axis shows the HSP90 binding score (log_2_ normalized to the wild-type BRCT).^31^ The z-axis heatmap shows the predicted viability of each variant as compared to wild-type cells (blue, low survival and weak resistance; red, medium survival and partial resistance; cyan, high survival and strong resistance). The connecting contours were generated by cubic interpolation using Python (see Methods). (F – G) The position of HSP90-buffered sites in the BRCT structure and (G) residue solvent accessibility. Residues exhibiting the strongest response to HSP90 inhibition are shown as red spheres (n=21). The number of variants in each bin is shown in parenthesis above the stacked bar. Statistical significance was determined using multiple two-tailed Mann-Whitney *t*-tests with multiple corrections applied (A and C). \*\*\*\**p*≤0.0001. \*\*\**p*≤0.001. \*\**p*≤0.01. \**p*≤0.05. ns, *p*>0.05. Data are presented as mean ± standard deviation values from three independent experiments.

To test if HSP90 prevents the proteolytic degradation of BRCA1 variants in cells, we measured mutant protein turnover in Cal 27 cells treated with HSP90 inhibitors. When we blocked new protein synthesis using cycloheximide (CHX), the moderately-engaged variant A1708V was turned over faster than the wild-type BRCA1 (Figure 3B). Indeed, six hours of low-level HSP90 inhibition depleted the A1708V variant similar to the severely destabilized S1715R variant. In addition, the wild-type BRCA1 protein was significantly more stable than A1708V despite some turnover, which is consistent with previous reports suggesting the wild-type BRCA1 requires HSP90 for stability.^71^ Furthermore, the HSP90-dependence of A1708V stability was reproduced using STA-9090 (Figure S3D). In contrast, we observed minor differences in turnover between A1708V and wild-type when HSP90 was not inhibited (Figure S3E). The severely destabilized S1715R variant exhibited markedly reduced expression and rapid turnover under all conditions.

To validate these results, we next tested the effect of heat stress on mutant cells that depend on HSP90 which we previously showed can inhibit the HSP90-buffering of *FANCA* mutations.^46^ In line with the increased environmental sensitivity of HSP90-engaged variants, the BRCA1-A1708V protein exhibited an accelerated rate of turnover at 39 °C as compared to the wild-type (Figure S3F). Such febrile-range heat stress significantly reduced the viability of *BRCA1* mutant Cal 27 cells only for the HSP90-engaged variants (Figure 3C). A1708V exhibited a significant decrease in cell survival of 31% as compared to the wild-type which was not affected. We reproduced this finding in HEK293A cells (Figure S3G). The effects were specific to the BRCA1 variant expressed because when dTAG treatment was not added there was no significant effect in cell survival (Figure S3H). In addition, we showed that mutant-specific loss of viability was not due to differences in proteostasis induced by heat stress on the different BRCA1 mutant cell lines, as determined using an assay to detect protein aggregates in cells.^72^ Indeed, wild-type, A1708V, and S1715R expressing mutant Cal 27 cells exhibited no difference in bulk protein aggregation at 37 °C or 39 °C, even after treatment with the proteasome inhibitor MG132 (Figure S3I). We conclude that HSP90 directly stabilizes BRCA1 mutants within cells and low-level HSP90 inhibition uncovers HSP90-buffered BRCA1 mutations by destabilizing the encoded protein variants.

Our results identify four criteria required for the HSP90-buffering of BRCA1 variants. These mutant proteins 1) bind moderately to HSP90 and not strongly to HSP70,^31,46^ 2) retain full or partial binding to partners, 3) support cell survival, and 4) exhibit increased turnover rate upon HSP90 inhibition. To better predict HSP90 buffering based on domain structure, we asked if HSP90-buffered BRCA1-BRCT variants share structural commonalities. Because it is not feasible to evaluate all four criteria for every clinically relevant BRCA1-BRCT variant, we developed a naïve approach to predict mutant cell survival based on a previously reported ML-based algorithms used to evaluate the dependence of alpha-1-antitrypsin variants on chaperone GRP94.^73^ We used the viability for known variants weighted based on the magnitude of HSP90 / HSP70 chaperone binding which correlates with mutant cell phenotype^31^ (Figure S3J and Methods). Projecting the predicted mutant cell survival onto a contour map revealed that HSP90-buffered BRCA1-BRCT variants occur throughout the domain (Figures 3D, 3E, and S3K). Indeed, structure mapping residues that result in the greatest sensitivity to HSP90 inhibition revealed an abundance of HSP90-buffered sites throughout the BRCT domain (Figures 3F). In addition, we observed a significant enrichment of HSP90-buffered variants for complex variants that partially disrupt folding and partner binding (Figure S3L). Notably, residues harboring HSP90-buffered sites tended to be more frequently buried inside the core of the BRCT domain (Figures 3G and S3M). These results suggest that HSP90 can buffer a plethora of BRCA1- BRCT domain mutations in humans, suggesting their clinical relevance.

### HSP90 buffering modifies the clinical significance of an abundant BRCA1 variant class

In light of our finding that HSP90 can buffer the deleterious effects of BRCA1 variants on cell viability, we next tested if the HSP90-buffering of BRCA1 variants can promote therapy resistance to genotoxic agents in cancer cells. Indeed, results from DNA repair and genome instability functional assays^58^ showed that HSP90-buffered BRCA1 variants frequently retained function (Figure S4A). To validate this observation, we tested the ability of three HSP90-buffered BRCA1 variants—A1708V, L1705R, and Y1845D—to promote resistance to PARP inhibition in cancer cells. Each of the variants independently supported cell survival when treated with PARP inhibitor olaparib in Cal 27 cells (Figure 4A, A1708V: 73 ± 5.3%). In contrast, S1715R and BRCA1-null cells were hypersensitive to low concentrations of olaparib, consistent with the synthetic lethality relationship between BRCA1-null cells and PARP inhibitors.^74–76^ Moreover, we validated this result in two different *BRCA1^dTAG/dTAG^*clones (Figures 4A and S4B). At the highest concentration tested, olaparib sensitized HSP90- buffered variant L1705R, but the wild-type cells also showed reduced survival under these conditions (Figures 4B and S4C). These results suggest that HSP90-buffered BRCA1 variants can promote therapy resistance in cancer cells.

**Figure 4.**
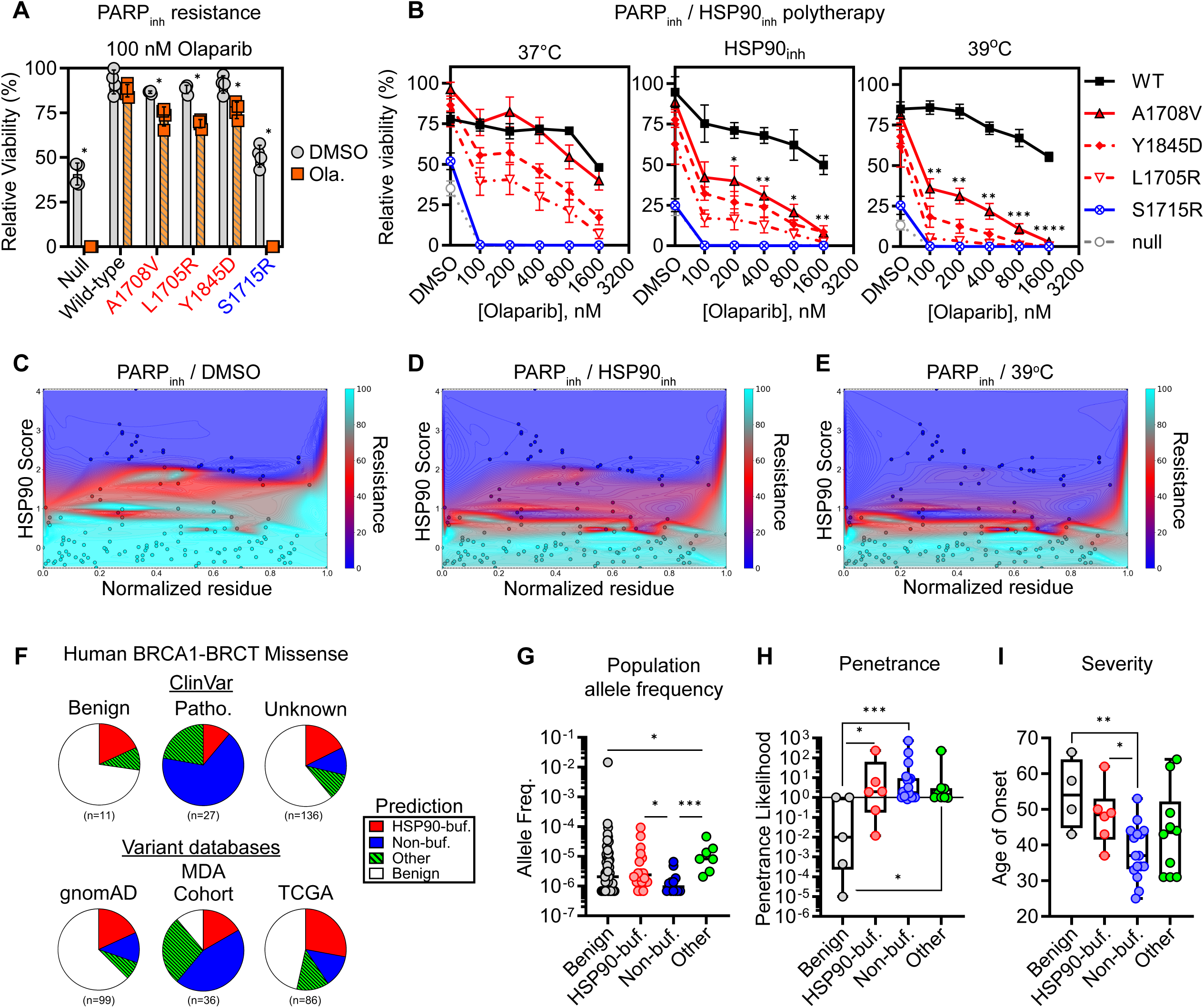
HSP90 buffering modifies the clinical significance of an abundant BRCA1 variant class. (A) Cell survival of Cal 27 cancer cells expressing HSP90-buffered BRCA1 variants (red). To ensure PARP inhibitor treatment was effective, these experiments leveraged high cell seeding densities where BRCA1-null cells retain partial survival under basal conditions. Indeed, BRCA1-null and S1715R mutant cells treated with 100 nM olaparib exhibited dramatically reduced cell survival. Cell survival measured by crystal violet. Ola., olaparib. (B) HSP90_inh_ / PARP_inh_ polytherapy specifically sensitizes three resistant HSP90-buffered cancer cells. HSP90 inhibitor was 15 nM NVP-HSP990. (C – E) Contour maps illustrating BRCA1-BRCT variant position, HSP90 binding score, and the effect of PARP / HSP90 inhibitor polytherapy on cell survival. Panel (C) shows 400 nM olaparib treated cells, panel (D) shows olaparib / 15 nM NVP-HSP990, and (E) shows olaparib / 39 °C co-treatment. (F) HSP90-buffered variant percentages for BRCA1-BRCT missense mutations observed in humans. Buffered variants were defined as moderately-engaged by HSP90 that do not mutate pSXXF coordinating residues (see Methods). Non-buffered variants comprise mutants strongly-engaged by HSP90 which are prone to severe folding defects. The ‘other’ bin includes splice site variants,^82^ contact variants,^69^ complex contact / folding variants, and other loss of function variants previously reported by comparing chaperone binding with deep mutational scans.^3,31^ The number of variants represented in each pie shown underneath as n. (G – I) The clinical significance of HSP90-buffered BRCA1-BRCT missense variants. (G) The population allele frequencies^85^ for different variant classes. (H) Penetrance likelihood^86^ and (I) clinical severity of HSP90-buffered alleles carried in humans.^115,116^ Statistical significance was determined using multiple two-tailed Mann-Whitney *t*-tests with multiple corrections applied (A, B, G, H, I). \*\*\*\**p*≤0.0001. \*\*\**p*≤0.001. \*\**p*≤0.01. \**p*≤0.05. All cell survival measurements were performed in 96- well plates at high seeding density to retain partial survival of null cells in the DMSO condition. Data are presented as mean ± standard deviation (A) and standard error of the mean (B) values from at least two independent experiments.

To determine the role of HSP90 in supporting the residual olaparib resistance of cancer cells expressing HSP90-buffered *BRCA1* mutations, we defined a range of HSP90 inhibitor concentrations that are nontoxic to wild-type cells but impact the survival of mutants to olaparib. For this, we compared the olaparib sensitivity of A1708V mutant cells and wild-type cells at progressively increasing doses of the HSP90 inhibitor NVP-HSP990. We observed that 10 nM NVP-HSP990 specifically reduced the cell survival of A1708V mutant cells to 39 ± 11% while wild-type cells retained 85 ± 9.1% viability at 800 nM olaparib (Figures S4D and S4E). Using a concentration of 15 nM NVP- HSP990, we found that HSP90-buffered mutant cells were more sensitive to olaparib as compared to wild-type cells which were not significantly affected compared to olaparib alone (Figure 4B). Moreover, exposure at febrile-range temperatures for the duration of the treatment hypersensitized the HSP90-buffered mutants to olaparib without affecting the wild-type (Figure 4B), demonstrating the increased sensitivity of the HSP90-buffered BRCA1 variants to HSP90 stress. These results suggest that low dose HSP90 / PARP inhibition combination polytherapy may be a viable approach to target HSP90-buffered BRCA1 mutations in tumors.

We re-applied our cell viability extrapolations / interpolations again, this time using the magnitude of chaperone binding as a diagnostic of PARP inhibitor response. In this way, we were able to broadly evaluate the efficiency of our HSP90 / PARP polytherapy for variants throughout the BRCT domain. Cancer cells with HSP90-engaged BRCA1 variants were predicted to resist PARP inhibitor therapies more than strongly HSP90-bound variants that were not buffered (Figure 4C). Moreover, the combination of PARP and HSP90 inhibitors specifically impacted the function of the HSP90-buffered variants (Figure 4D). In contrast, wild-type-like variants that weakly bound to HSP90 were predicted to remain unaffected by the combination therapy. The same assumption was used for cells co-treated with febrile-range heat shock, given that at 39 °C HSP90-buffered BRCA1 mutants lose function (Figure 4E). Under all three conditions, the C-terminal region of the domain was more resistant to mutation as evidenced by the upward curvature in the resistance landscape, consistent with prior reports.^77^ Importantly, we found that HSP90-buffered variants permeate the entirety of the BRCT domain.

Next, we asked whether machine learning algorithms can identify BRCA1-BRCT mutations we predicted to be buffered. Strikingly, the pathogenicity of HSP90-buffered variants could not be distinguished from loss-of-function mutations by four different next-generation machine learning approaches (Figures S4F-I).^78–81^ Only when consolidating known contact and other hypomorphic mutations into a separate bin (‘other’: splicing, pure contact, and complex contact/folding variants)^69,82^ did we observe a significant difference in the pathogenicity of HSP90-buffered and non-buffered alleles (Figure S4J). Nevertheless, prediction tools assigned highly variable pathogenicity scores to both HSP90-buffered and known hypomorphic *BRCA1* mutations (Figure S4J), suggesting there is room for improving the ability of these algorithms to accurately classify mutations based on their severity.

Our approach to classify HSP90-buffered *BRCA1* mutations enabled us to next examine the clinical manifestations of HSP90 buffering. HSP90-buffered *BRCA1* mutations were prevalent in ClinVar,^63^ as well as in three other population genomics datasets (Figure 4F).^31,83–85^ We estimate that 11-28% of known human BRCA1-BRCT missense variants are buffered by HSP90. These results highlight the importance of stratifying HSP90-buffered mutations in the general population.

Finally, we determined the clinical significance of HSP90-buffered *BRCA1* variations on cancer risk in human carriers. HSP90-buffered BRCA1 variants appear to accumulate in the general population similar to those we predicted to be benign (Figure 4G). However, inherited HSP90-buffered variants from the Leiden Open Variant Database (LOVD)^86^ were similarly penetrant to non-buffered high-penetrance *BRCA1* alleles (Figure 4H). These results suggest that HSP90 promotes the accumulation of *BRCA1* mutations in the population, at the cost of increasing the inherent risk of cancer development. Moreover, when investigating the clinical severity of HSP90-buffered mutations, we observed a ∼10 year delay in breast cancer onset for patients carrying HSP90-buffered alleles as compared to non-buffered alleles (Figure 4I). Hence, by buffering *BRCA1* mutations, HSP90 appears to delay cancer onset in humans. In conclusion, our data shows that HSP90 buffers deleterious genetic variations in *BRCA1*, profoundly impacting the clinical outcome of breast cancer patients carrying these variants.

## DISCUSSION

The protein-folding chaperone HSP90 buffers the phenotypic consequences of genetic and epigenetic variation in ecologically relevant settings.^39,40,43,45,87,88^ We previously suggested that HSP90 buffering explains the variable expressivity and environmental sensitivity of inherited mutations that cause the rare genetic disorder Fanconi anemia,^46^ but the scope and significance of this finding to common genetic disorders remained unknown. Here, we report that HSP90 buffers deleterious mutations in the most well-studied cancer predisposition gene, *BRCA1*,^89–96^ and in so doing, HSP90 modifies the PARP inhibitor sensitivity of cancer cells and age of cancer onset in humans. We defined the criteria of HSP90-buffered missense mutations and revealed their prevalence in the general population. This work demonstrates the broad significance of HSP90 buffering in cancer predisposition.

A class of variants in the BRCT domain of BRCA1 moderately bind HSP90, retain interactions with DNA repair complexes, and support cell survival, indicating that HSP90 salvages variant function. However, HSP90-buffered variants acquire sensitivity to environmental proteotoxic exposures that compromise HSP90 function within the cell. Indeed, the function of HSP90-buffered variants were hypersensitive to febrile-range heat shock as well as to nontoxic concentrations of small molecule HSP90 inhibitors. Moreover, HSP90-buffering of BRCA1 variants increased the resistance of cancer cells to PARP inhibitor overcome by supplementing low-level HSP90 inhibitor. Hence, identifying HSP90-buffered alleles is anticipated to be critical for improving the efficacy of PARP inhibitors by stratifying patients for combination therapies based on low-level HSP90 inhibition. In breast cancer patients, HSP90 buffering delayed the age of disease onset by ∼10 years, indicating that HSP90 can alter the expressivity of heritable breast cancer. We estimate that ∼11-28% of BRCA1-BRCT missense variants are HSP90-buffered and thus a substantial percentage of cancer patients may benefit from the combined targeting of BRCA1 and HSP90.

In regards to the structural features of HSP90 buffering, we identified HSP90-buffered variants throughout the BRCT domain of BRCA1, involving numerous secondary and tertiary structural elements. Most of the HSP90-buffered variants we identified were buried in the BRCT structure, suggesting that partial unfolding of the hydrophobic core can be stabilized by HSP90 chaperoning. These findings are consistent with recently determined HSP90-GR structures showing that GR acquires partially native structure.^97,98^ Interestingly, contact mutants on the domain surface previously described as hypomorphic also accumulate in the population and confer milder impact on cancer risk similar to HSP90-buffered variants.^99–101^ Even the complex contact / folding variant R1699W, which disrupts both pSXXF binding and domain folding, is predicted to be at least in part buffered by HSP90. Indeed, in yeast, the function of R1699W is temperature sensitive.^102,103^ The temperature sensitivity of HSP90-buffered variants suggests the structure is more metastable than the wild-type. HSP90 binding to the partially folded conformations of HSP90-buffered BRCA1 variants acts to salvage them within the cell. Predicting the inherent metastability of mutant proteins could enable tailoring disease diagnosis, prognosis, and management to the environmental sensitivities this metastability confers. Whether inherent metastability of proteins variants is common in other structured domains remains unknown, substantiating the screening for HSP90-buffered variants in other disease risk genes.

HSP90 buffering has important implications for cancer therapy. HSP90 inhibition monotherapies were not sufficient to completely disrupt *BRCA1* mutant cells carrying HSP90-buffered alleles, and increasing HSP90 inhibitor doses induced general toxicity to normal cells. These results are consistent with the toxicity observed in clinical trials that use HSP90 inhibitors.^70,104–107^ On the other hand, HSP90-buffered *BRCA1* variants confer full to partial resistance to PARP inhibitor olaparib in cancer cells under basal conditions. Our results suggest that HSP90 buffering can account for much of the unexplained resistance to PARP inhibitors observed in 40% of cancer patients with *BRCA1* mutations.^75,91,92,96,108–110^ The data also provide a framework to stratify patients expected to respond well to HSP90 inhibitor combination therapies.^111,112^ Indeed, low-level HSP90 inhibitor concentrations hypersensitized cancer cells expressing HSP90-buffered BRCA1 variants to PARP inhibition. In this work, we demonstrate that HSP90 buffers a predominant class of deleterious genetic variations in humans, and thus, alters the risk of cancer development.

### Limitations of the Study

Our study thoroughly investigated a significant number of clinically relevant mutations in one domain of interest, identifying a small number of mutation profiles that could infer HSP90 buffering. Although no general principles linking HSP90 buffering and specific structured elements or motifs emerged, such associations may exist, as HSP90-buffered variants may be more or less abundant in certain domains. In addition, our study specifically investigated the role of HSP90 buffering in PARP synthetic lethality, but platinum-based drugs are also of interest since they are commonly administered for cancer therapy. Another limitation of this study is that it does not examine clinically relevant environmental factors that may reveal HSP90-buffered BRCA1 mutations in humans. Furthermore, *ex vivo* results do not always recapitulate the effects of inhibitors *in vivo*. It remains to be tested if HSP90 inhibitors can specifically sensitize HSP90-buffered mutant tumor cells to therapies *in vivo*. Moreover, HSP90 may impact the function of diverse mutation types and drive therapy resistance, including frameshifting variants as previously described.^75,92^ Regardless, our discovery of HSP90-buffered mutations in *BRCA1* warrants experiments to test HSP90’s role in therapy resistance in mouse tumor models and the design of clinical trials to develop its applications in precision medicine.

## ACKNOWLEDGEMENTS

We thank John Tainer, Richard Wood, Swathi Arur, Adriana Paulucci, Richard Behringer for critical feedback, and the MD Anderson Editing Services Research Medical Library on editing the manuscript. We thank Natalia Condic, Beryl John, Xing-Han Zhang, and Tin Pham for assistance with CRISPR/Cas9 editing, flow cytometry, and cell sorting. We thank Adriana Paulucci-Holthauzen for assistance with training, troubleshooting, and maintenance of high content microscope imaging and analysis in the Genetics Department BSRB Microscopy Core at MD Anderson. We thank Gregorz Sienski for guidance on CRISPR/Cas9 genome engineering at the *HBEGF* locus. We thank Pam Rowling and Laura Itzhaki for providing quantitative pSXXF binding data and Bin Wang for guidance and for providing homemade Abraxas antibody. We acknowledge support from the Cytometry and Cell Sorting Core at Baylor College of Medicine with funding from the CPRIT Core Facility Support Award (CPRIT-RP180672) and the NIH (CA125123 and RR024574), and with the assistance of J. M. Sederstrom, and the Flow Cytometry & Cellular Imaging Core Facility, which is supported in part by the National Institutes of Health through M. D. Anderson’s Cancer Center Support Grant P30 CA016672, and with the assistance of N. R. Vaughn. This research was supported by the National Cancer Institute of the NIH under Awards F32CA253780 (B.G.) and K22CA222938 (G.I.K.). The content is solely the responsibility of the authors and does not necessarily represent the official views of the National Institutes of Health. G.I.K. is a Cancer Prevention & Research Institute of Texas (CPRIT) Scholar in Cancer Research. This work was supported by CPRIT grant RR180005 (G.I.K.), a Kleberg Innovator Award (G.I.K.) by the Robert J. Kleberg, Jr. and Helen C. Kleberg Foundation, and a Basser External Research Grant by the Basser Center for BRCA (G.I.K.).

## AUTHOR CONTRIBUTIONS

B.G. and G.I.K. conceived the project. B.G. designed and performed experiments with assistance from P.M. M.H. and B.G. performed degron-tag cell line engineering using CRISPR/Cas9 with assistance from J.C. B.G. and P.M. performed CRISPR/Cas9 genome engineering and P.M. performed quality control FACS and flow cytometry assays. B.G. and G.I.K. wrote the manuscript with input from all co-authors.

## DECLARATION OF INTERESTS

The authors declare no competing interests.

**STAR★METHODS**

## EXPERIMENTAL MODEL AND SUBJECT PARTICIPANT DETAILS

### Mammalian cells

HEK293A and Cal 27 cells were propagated in 10-cm petri dishes (Corning, 08-772-22) at 37°C with 5% CO_2_ in Dulbecco’s modified Eagle’s medium (Corning, 10-013-CV) supplemented with 10% fetal bovine serum (Sigma-Aldrich, F2442) and 1% Pen-Strep (Gibco, 15140-122). Once cells reached ∼70% confluency they were washed once with PBS (Gibco, 10010023), treated with Accumax (Innovative Cell Technologies, AM-105), incubated at 37°C for 15-20 min, and diluted in normal media to passage cells to a new petri dish at a 1:10 dilution. All cell lines were validated using STR fingerprinting at the MD Anderson Cytogenetics and Cell Authentication Core and tested negative for Mycoplasma contamination (Lonza, LT07-318). Cell lines sorted using fluorescent reporter proteins were evaluated for stable fluorescent expression every 5-10 passages and all experiments were performed using cells passaged up to 25 times after sorting.

BRCA1 variants from human mutation databases ClinVar, TCGA, and gnomAD were previously described.^31^ BRCA1 mutant allele frequency data was taken from version 4.1.0 of the gnomAD database.^85^ BRCA1 mutant penetrance data and human breast cancer patient age of onset was taken from our previously described work.^31^

## METHOD DETAILS

### Plasmids and genome engineering

Dual expression plasmid pUX459 Cas9/gRNA targeting the endogenous *BRCA1* and pUC19 degron-tag repair template (BSD-2A-2xHA-FKBP12^F36V^) were used for degron tag knock-in at the N-terminus of the endogenous *BRCA1* locus. Plasmids in the pVAX1 backbone expressing Cas9 or gRNA targeting the *HBEGF* gene locus and pUC19 BRCA1 repair template were described previously.^57^ Unless otherwise noted, all BRCA1 engineering constructs included a HBB IVS2 intron sequence 3′ of the mRuby3 sequence (BRCA1-2A-mRuby3-intron) which improved the long-term stability of the knock-in. BRCA1 mutant plasmids were cloned using oligonucleotide primers containing the mutation (Sigma-Aldrich) and two-step stitch PCR to generate double-stranded DNA products harboring the mutation. PCR products were purified by agarose gel extraction (Takara Bio, 740609) and the wild-type plasmid was linearized using restriction enzymes (New England BioLabs). In-Fusion® cloning (Takara Bio, 638918) was then used to assemble the linearized plasmid and mutant PCR products. Full-length N-terminally 3xFLAG tagged BRCA1 was cloned by the same approach using a β-Globin-3xMyc-BRCA1 plasmid digested to replace the 3xMyc tag with 3xFLAG. BRCA1 truncation 1,314- 1,863 fused to N- or C-terminal 3x-FLAG tag were previously reported.^31^

To generate monoclonally-derived *BRCA1^dTAG/dTAG^* HEK293A and Cal 27 homozygous knock-ins, we used BRCA1 repair donor templates that included a blasticidin resistance gene and a 2A ribosomal skipping sequence. Cells engineered in-frame acquire blasticidin resistance, enabling selection of knock-ins before monoclonal derivation. Cells were collected and counted with Trypan Blue using a Countess II (Life Technologies) and diluted to seed 1 million cells in 10-cm dishes at Day 0. On day 1, cells were forwarded transfected by mixing 9000 ng of pUX459 Cas9/gRNA with 9000 ng of pUC19 BRCA1 repair templates and packaged using 2400 µL of OptiMEM combined with 48 µL of Lipofectamine2000 following incubations as recommended by the manufacturer (Invitrogen, 11668). On day 2, transfection media was removed and replaced with fresh warm media and on day 4, Blasticidin (Life Technologies, A1113903) was added to a final concentration of 5 µg/mL, and Blasticidin was replenished for a second treatment on day 7. Following Blasticidin treatment on every subsequent third day, the media was replenished with fresh warm media. On day 18, genomic DNA was purified using DNAeasy blood and tissue kit (Qiagen, 69504) and the knock-in efficiency was evaluated by PCR of the 5’ region of the *BRCA1* locus. Successful integration of the dTAG knock-in resulted in a distinct ∼2.4 kb PCR product in contrast to the parental cells which exclusively exhibited a ∼1.55 kb product consistent with the sequence of the endogenous *BRCA1* locus. Moreover, PCR using a FKBP12-specific primer set yielded a ∼1.2 kB PCR product only for monoclonally derived knock-in cells because parental cells did not include a priming site for the FKBP12-specific primer. Following PCR-based validation of the knock-in, cells were lifted, counted, and diluted to a concentration of 0.6 cells per 100 µL and seeded into ten 96-well plates (Corning, 3598) at 100 µL per well. Individual seeded cells were identified using brightfield microscopy and monoclones were grown for 16 days followed by expansion in 6-well plates (Corning, 3506) to collect genomic DNA for PCR screening. Monoclonally derived homozygous knock-in cells were identified using the same PCR-based approach.

For genome engineering at the *HBEGF* gene locus, 50,000 HEK293A cells were reverse transfected using 60 ng each of Cas9 expressing plasmid, *HBEGF* intron 3 gRNA plasmid, and 3xF-BRCA1-2A- mRuby3 repair template plasmid in 50 μL of PEI MAX® (Polysciences, 24765) with OptiMEM (Gibco, 319850) at a ratio of 1:100 (10 μg/mL final) using incubation times according to the manufacturer. The next day, cells were collected and re-seeded in 6-well plates and grown for 2 days after which media was supplemented with 20 ng/mL diphtheria toxin (DT).^57^ Cells successfully engineered acquire resistance to DT, generating a polyclonal pool of fluorescent cells suitable for purification by FACS. Fresh warm media was exchanged twice every 3 days followed by expansion in 10-cm dishes before sorting by FACS. Cal 27 cells were engineered in the same fashion except 200,000 cells were seeded in 6-well plates and forwarded transfected with 1000 ng of each plasmid using Lipofectamine2000. Sorting was carried out on a BD FACSAria II cell sorter with a 1.0 OD neutral density filter using the following laser configurations for mRuby3: 561nm EX; 610/20nm EM. After gating for single cells, 10,000 single cell events were recorded for each cell line. Sort gates were then drawn such that the brightest 50% of each fluorescent population was sorted. For each cell line, the entire sample was sorted. The total number of cells collected ranged from 400,000-4,000,000 cells per cell line.

To confirm the genotype at the endogenous *BRCA1* and *HBEGF* gene loci after sorting, gDNA specific primers were designed to amplify each gene. For detecting *BRCA1^dtag/dtag^*, the N-terminal region of BRCA1 was PCR amplified and PCR products gel extracted. Premium PCR sequencing was carried out by Plasmidsaurus. For detecting *HBEGF^BRCA^*^1*/BRCA1*^ mutants, the C-terminal region of BRCA1 was PCR amplified, gel extracted, and sequenced by Sanger Sequencing at Epoch Life Science. To measure cell growth and viability, an aliquot of cells were stained and counted using Trypan Blue.

### CHX chase, Co-IP, dTAG degradation, and Western blotting

For CHX chase experiments, Cal 27 cells stably expressing 3xFLAG-BRCA1 variants were seeded in 6-well plates at 15,000 cells per well. After 2 days, cells were treated with 100 μg/mL CHX (Sigma-Aldrich, C7698) plus or minus HSP90 inhibitor in a staggered-way so that cells were collected at the same time. Cells were PBS washed and treated with accumax to lift and collect into 1.5 mL Eppendorf tubes. Samples were centrifuged at 500xg for 5 min, accumax aspirated, and cells directly lysed in urea/SDS protein sample buffer followed by denaturation for 5 min at 65°C. This protocol improved reproducibility and evenness for protein turnover experiments.

CoIP was performed by reverse transfecting 220,000 HEK293T cells with 875 ng of plasmid in 24- well plates (Corning, 3524). After 2 days, cells were place on ice, washed once with PBS, and incubated with lysis buffer for 15 min (50 mM HEPES-KOH, pH 8.0, 150 mM NaCl, 10 mM MgCl2, 20 mM NaMo, 0.7% Triton X-100, 5% glycerol) supplemented with protease inhibitors (Leupeptin, Sigma-Aldrich, 11034626001; Aprotinin, Santa Cruz Biotechnology, sc3595; Pepstatin A, VWR, 97064-248; PMSF, Sigma-Aldrich, 7110), phosphatase inhibitors (Sodium Orthovanadate, New England BioLabs, P0758; Sodium Fluoride, Sigma-Aldrich, S7920), RNAse A (Fisher Scientific, EN0531) and Benzonase (MilliporeSigma, 70664). The soluble fraction was collected by centrifugation at 14,000xg for 10 min at 4°C. The soluble lysate was then mixed with 10 μL of pre-washed agarose-linked anti-FLAG beads (Sigma-Aldrich, F2426) and incubated for 3 h at 4°C with gently mixing. Non-specific interactions were diluted by washing four times with ice-cold lysis buffer and flag-tagged proteins were eluted using 150 μg of 3xFLAG peptide (Biomatik, 56305) for 30 min at room temperature. Immunoprecipitated and eluted flag-tagged proteins were denatured using an equal volume of urea/SDS protein sample buffer followed by incubation at 65°C for 5 min.

To measure dTAG-BRCA1 degradation using dTAG^V^-1 compound (Tocris Bioscience, 6914) as compared to dTAG^V^-1-NEG control (Tocris Bioscience, 6915), 25,000 cells were seeded in 24-well plates and 2 days later dTAG treatments were administered in a staggered fashion for indicated times. Cells were lysed and collected using the same lysis buffer as for coIP. Whole cell extracts were mixed with an equal volume of urea/SDS protein sample buffer and denatured at 65°C for 5 min.

Proteins were resolved by molecular weight using pre-cast 4-12% SDS-PAGE gels (Invitrogen, WG1402) run for 4 hours at 160V on ice in standard SDS-MOPS buffer (Invitrogen, NP0001). Proteins were then transferred to PVDF membranes (Bio-Rad Laboratories, 1620177) using wet transfer in tris-glycine buffer with 20% methanol by running for 3 hours at 0.3 constant amps at 4°C. Membranes were blocked for 60 min at room temperature using 5% milk (Lab Scientific, M0841) in PBST (0.05% Tween 20) followed by incubation with primary antibodies overnight at 4°C. The Abraxas antibody was a gift from Bin Wang. To visualize proteins, membranes were washed thrice in PBST for 5 min each and the incubated with anti-mouse (Sigma-Aldrich, F1804) or anti-rabbit (Sigma-Aldrich, A0545) peroxidase secondary antibodies prepared in 5% milk for 60 min. Membranes were washed again twice with PBST and once with PBS followed by 1-2 min incubation in horseradish peroxidase substrate (Millipore, WBKLS0) before visualization using a Chemidoc MP imaging system (Bio-Rad Laboratories).

### Crystal violet and CellTiter-Glo® assays

On day 0, cells were collected by accumax treatment, re-suspended in cell culture media, and placed to ice for counting and calculating dilutions. Cells were diluted and seeded in 0.5 mL in 24-well (crystal violet staining) or 0.1 mL in 96-well plates (crystal violet staining and CellTiter-Glo®). 96-well plates were seeded with cells using an automated liquid handling robot (Biotek, EL406). Experiments in 24-well plates were performed with technical triplicates and 96-well plates were performed in technical quadruplicates or duplicates. After seeding, cells were allowed to settle at room temperature for 30 min and then put to incubate at 37°C with 5% CO_2_.

On day 1, cells were treated with compounds by preparing 5-fold (24-well) or 3-fold (96-well) concentrated dilutions and adding 0.125 mL (24-well) or 0.05 mL (96-well) directly to plates. Unless otherwise stated, dTAG concentrations were 0.5 µM. Inhibitor titrations were performed in 96-well plate formats using multi-channel to generate serial dilutions and add drug treatments. Cells to be heat shocked were then placed at 39°C with 5% CO_2_ at this time.

Both high and low seeding densities were tested to determine optimal conditions for measuring BRCA1 variant function in cell survival. High and low seeding in 24-well plates corresponded with 1,000 and 250 cells, while in 96-well plates it was 400 and 100 / 25 cells seeded. At high cell seeding, the effect of BRCA1 depletion on cell survival was dampened presumably because loss of BRCA1 results in a defect in cells growth rate at early time points on the time-scale of our experiments.^3^ In contrast, HSP90 or PARP inhibitor treatments resulted in clear cell killing effects as observed by small round cellular morphology and detachment from the culture plate.^74^ Hence, for PARP inhibition experiments high cell seeding densities were utilized because the effect on cell survival was observed to be highly robust. Cells were incubated for 6-7 days at high cell seeding density or 9-10 days at low cell seeding density which were shown to be within the linear range of the cell survival assay.

Cells were terminated either by staining with crystal violet or treatment with CellTiter-Glo® reagent. Staining cells with crystal violet was carried out as previously described.^117^ Culture media was manually aspirated followed by incubation with 100% methanol for 30 min. Cells were then washed once with deionized water and then incubated in crystal violet staining solution for 60 min (25% methanol and 0.5% crystal violet powder (Sigma, C6158). Plates were washed under running water 4 times and left to air dry overnight before imaging. For 96-well plates, media was aspirated and cells were washed thrice with PBS using an automated liquid handling robot (Biotek, EL406). 0.2 mL of crystal violet solution was then added using a multi-channel pipet treated the same as 24-well plates. CellTiter-Glo® was carried out as recommended by the manufacturer with some modifications. Plates were incubated on benchtop for 15 min to equilibrate to room temperature. CellTiter-Glo® reagent was dispensed using a multi-channel pipet at 1:5 dilution factor followed by spin down at 300xg for 1 min. Plates were incubated on a plate rocker for 5 min and luminescence read using a 400 to 700-nm filter equipped to an EnVision Plate Reader (PerkinElmer).

### Aggregate detection assay

Aggregates within cells were detected using a commercially available kit as recommended by the manufacturer (Enzo Life Sciences, ENZ-51035). On day 0, 50,000 Cal 27 cells were seeded in 6-well plates. On day 3, cells were treated with 10 µM MG132 and 0.5 µM dTAG for 16 hrs. Cells were then collected, fixed, permeabilized, and incubated with anti-Aggregate detecting antibody as recommended by the manufacturer. A Beckman CytoFLEX flow cytometer equipped with fluorescence filter set 561nm EX; 638nm EM was used to detect changes in aggregate abundance under different conditions. Aggregation propensity factor (APF) was calculated by normalizing the mean fluorescence intensity between two conditions being compared.

### BRCA1 variant predictions

Variants were selected from ClinVar, gnomAD, TCGA/cBioPortal, LOVD, and a prospective MDA breast cancer patient cohort as previously described.^31^ Collected BRCA1-BRCT variants assays were mined from NeXtProt server which classifies the effect of mutation as a phenotypic intensity for each assay.^58^ To determine if variants that moderately engaged HSP90 exhibit partial functional activity, the number of assays reported as “mild” or “moderate” effect for each variant was calculated and expressed as a percentage relative to the total number of assays reported for each individual variant. Hence, variants that were frequently functional or severe phenotype intensity exhibit low percent assays partial function. Only variants that reported 3 or more assays were considered. The percentage of assays exhibiting partial function were then binned by HSP90 interaction magnitude or pSXXF disrupting. pSXXF disrupting variants were previously classified by comparing variant chaperone binding, cell survival, phosphopeptide binding, and proximity to the pSXXF in the BRCT crystal structure.^3,31,60,61,118^ Similarly, the effect of BRCA1-BRCT mutation on pSXXF binding was determined by considering previously reported results.^31,61,62^ First, known pSXXF disrupting variants were assigned as “severe” variant PPI effect. Then, variant pSXXF binding activity and specificity were considered. Variants that exhibited severe effects in activity or specificity were assigned “severe”, variants that exhibited partial activity in binding or activity were assigned “partial”, and variants that retained pSXXF binding activity and specificity were assigned “no effect”.

We took inspiration from a machine learning based approach use to predict the effect of genetic variation and GRP94 on the function alpha-1-antitrypsin protein.^73^ BRCA1 variant cell survival extrapolations / interpolations were calculated using a weighted and vectorized chaperone interaction approach based on measured PARP inhibitor response. This approach leverages the fact that the magnitude of HSP90 / HSP70 binding to BRCA1-BRCT variants is proportional to the functional defect of *BRCA1* mutations.^31^ First, we utilized our measured cell viability data on six BRCA1-BRCT variants (wild-type, A1708V, L1705R, Y1845D, S1715R, and A1708E), and HSP90 / HSP70 chaperone interaction scores for a collection of ∼140 variants, excluding pure contact mutants which do not correlated with the magnitude of chaperone binding. For variants that we did not measure the effect of mutation viability, the two nearest neighbors in chaperone interaction space with known viability response were identified by calculating the length of the vector between variants in HSP90 / HSP70 interaction space. The predicted survival was then calculated as the weighted viability effect based on the distance to the measured variant (e.g., a variant 0.5 interaction distance from a 50% viability variant and 0.25 interaction distance from a 100% viability variant is predicted to be 83.333% viable). Because the A1708E variant was not extensively studied due to anomalous gDNA PCR profile, we used all our available A1708E viability data and correlated it against S1715R viability data collected in the same experiment. The matched data were fit to a linear regression and unknown A1708E data was calculated from known S1715R data. To generate continuous viability prediction landscapes, a cubic interpolation was used in Python to generate a smoothened landscape with contours connecting the predicted variant viability data. For this, the maximum and minimum predicted mutant cell viability was set to 100% and 0%, respectively. In addition, four pseudo viability predictions were generated at each corner of the landscape corresponding to variants that N- and C- termini that weakly or strongly bind to chaperones.

To calculate HSP90-buffered sites from the prediction viability response data, the effect of HSP90 inhibitor NVP-HSP990 alone was compared with DMSO treated cells. First, we filtered out variants that exhibited viability <25% in DMSO treated conditions, as they variants are considered severely disrupted loss of function and do not fit the criteria of HSP90-buffered variants. Then, duplicates were removed for residues with more than one mutation in our data set, retaining only the mutation that resulted in the largest change in viability upon HSP90 inhibition. The distribution of these data were then considered and HSP90-buffered sites were assigned as those in the highest fourth quartile effect. The features of HSP90-buffered sites were then determined using the GetArea webserver^119^ on BRCT without the pSXXF present (accessibility), the STRIDE web portal^120^ (secondary structure elements), and the ProteinTools webserver^121^ (BRCT region). BRCT residue regions were uniquely assigned using a hierarchical assignment approach. First, pSXXF proximal variants were assigned as previously described.^31^ Then, long-range interactions were defined as residues within 5Å in the BRCT structure and 50 amino acids apart in primary sequence. Remaining residues part of BRCT1 and BRCT2 subdomains were assigned to the remaining residues with separation at residue 1,756.

HSP90-buffered, non-buffered, benign, and other variants were predicted based on HSP90 / HSP70 chaperone interaction magnitude, comparative functional data with cell survival in HAP1 cells, and established complex BRCA1-BRCT variants.^3,31^ Variants classified as ‘other’ included pSXXF disrupting variants, splice variants, and other variants that do not bind chaperones yet lose function in cell survival assays. Then, variants that bind strongly to HSP90, HSP70, or both chaperones were assigned as ‘non-buffered’. Variants that bound moderately to HSP90, HSP70, or both chaperones were predicted to be “buffered”. Lastly, remaining variants that weakly bind chaperones and do not disrupt pSXXF were predicted to be ‘benign’.

## QUANTIFICATION AND STATISTICAL ANALYSIS

### Quantifying relative cell viability

Crystal violet-stained plates were imaged with a High Content Analysis System composed of a Nikon Inverted Eclipse TiE microscope equipped with a 16.5 uM pixel DS Ri2 color camera using an 2X 0.10 NA Plan Apo Objective. A job automated acquisition was programmed to screen 24-well and 96- well plates, where a central region of interest (1,300 by 1,300 pixels) was imaged. For 24-well plates, each well was split into a 3×3 grid and stitched together by image alignment because the well could not be captured with one field of view. White balance and shade corrections were performed before acquiring each plate. Quantification of stained cells in each image was performed using pixel counting with saturation, hue, and intensity thresholding set to specifically capture stained cells. The sum density pixels were then counted within a region of interest using the automated Nikon acquisition package to determine the raw cell count of each well.

To convert raw sum density pixel counts to relative viability (%), the raw sum density of dTAG^V^-1 treated wells was normalized to the average of dTAG^V^-1-NEG treated wells. Averages were taken for each cell line separately to normalize and account for differences in cell seeding. This approach enabled the determination of cell survival using crystal violet staining under different conditions. To assess if cell survival measured using this approach reflected relative cell viability, the results were compared with cell viability as measured by CellTiter-Glo® assay. Cell viability was calculated using the same approach by normalizing luminescence signals for dTAG^V^-1 treated to the average of dTAG^V^-1-NEG treated wells. The correlation between these two approaches reached an R^2^ of 0.9, demonstrating that cell survival quantification by crystal violet staining and pixel counting also reflects relative cell viability. Unless otherwise noted, all viability data shows the average of independent biological replicate experiments carried out on different days and the associated standard deviation of the averages.

### Statistical analysis

Statistical analyses are described at the end of each figure legend. Analyses were performed using the GraphPad Prism software package. Significance was defined as p≤0.05 (indicated by a single *).

## SUPPLEMENTAL FIGURE TITLES

**Figure S1. *BRCA1^dTAG/dTAG^* knock-ins enable conditional complementation of *BRCA1* in multiple cell lines, Related to** Figure 1

(A, B) PCR-based approach to detect monoclonal cell lines carrying homozygous *BRCA1^dTAG/dTAG^* knock-in. Top panel in (A) shows PCR to amplify exon 2 of the endogenous *BRCA1* locus. Priming at the endogenous locus results in a PCR product of ∼1.55 kB compared to the knock-in product of ∼2.44 kB. The absence of the ∼1.55 kB band in the clone indicates a pure homozygous *BRCA1^dTAG/dTAG^* knock-in was engineered. Bottom panel in (A) shows PCR-based approach using one primer that targets the Fkbp12 region such that only genomic DNA carrying the knock-in sequence produces a product of ∼1.2 kB. (B) Sequence alignments against the endogenous *BRCA1* and knock-in sequences are shown as a solid arrow, with chromatograms for the intervening knock-in regions. The mutated gRNA binding site and PAM region are boxed. MW, molecular weight. kB, kilobase. L, ladder. P, parental cell line. C, monoclonally derived knock-in. CDS, coding DNA sequence. PA, Primer A. PB, Primer B. PC, Primer C.

(C) Cell counts and viability of parental cells and two knock-in Cal 27 clones as measured using Trypan blue staining. Cells were grown for three days and counted, then passaged at the same dilution and counted again three days later.

(D, E) Western blotting of monoclonally derived HEK293A and Cal 27 cell lines detected by anti-HA antibody (D) or anti-BRCA1 antibody (E). In (E), clones were treated with dTAG to test for conditional depletion of dTAG-BRCA1. HEK293A were treated with dTAG-NEG (NEG) or dTAG (+). P, parental. P+, parental treated with dTAG. MW, molecular weight. kDa, kiloDaltons. C1, C2,…, clone1, clone2.

(F, G) BRCA1 degradation time courses at short and long times points indicates conditional, rapid, and robust degraded of dTAG-BRCA1 in HEK293A and Cal 27 cells. Negative (–) and positive (+) controls were treated for 7 hours with dTAG and dTAG-NEG, respectively. HEK293A cells in (G) were treated with genotoxic agent MMC to demonstrate that DNA damage does not re-introduce endogenous BRCA1 expression (2+MMC). Cal 27 cells in (G) were grown to high density first and then treated with dTAG to demonstrate that dTAG is saturating for dTAG-BRCA1 degradation at low and high cell seeding (D6+).

(H – J) BRCA1 depletion using dTAG treatment decreases cell survival in a dose-dependent manner for HEK293A and Cal 27 cells in 24-well plate format. 2 µM dTAG treatment appeared toxic in Cal 27 cells. Hence, cells were treated with 0.5 µM dTAG in (J) for five monoclonally derived Cal 27 cell lines. Clone four was determined to not be homozygous knock-in (*) and exhibited insensitivity to the dTAG treatment.

(K) FACS to purify cells engineered at the *HBEGF* locus to express mRuby3 fluorescent reporter (left column). The vertical dashed line indicates the brightest fluorescent cells sorted. 3xF-BRCA1-2A- mRuby3 cells were reproducibly dimmer as compared to mRuby3 cells, likely owing to the larger BRCA1 expression sequence. Cell fluorescence was maintained after sorting for at least 2 months of passaging (right column). Negative control dark cells shown in cyan.

(L – M) Cell survival of HEK293A and Cal 27 cells complemented with 3xF-BRCA1 in 96-well plate format using crystal violet staining. 2 µM dTAG treatment was toxic in Cal 27 cells. BRCA1 complementation in clone 3 was reduced as compared to clone 2, so we proceeded with clone 2 as the primary Cal 27 *BRCA1^dTAG/dTAG^* cell line. Statistical significance was determined using two-tailed Mann-Whitney *t*-test (C) and Kruskal-Wallis ANOVA test (L). \*\*\**p*≤0.001. \*\**p*≤0.01. ns, p>0.05. Data are presented as mean ± standard deviation values from at least two independent experiments.

**Figure S2. Moderate HSP90 engagement signifies maintenance of BRCA1 function, Related to** Figure 2

(A) pSXXF binding affinity for *In vitro* purified BRCT variants^62^ correlated with the magnitude of HSP90 binding.^31^ Horizontal dashed lines show cutoffs for weak (HSP90 Score < 0.5), moderate (0.5 < HSP90 Score < 2.0), and strong (HSP90 > 2.0) binding. Vertical dashed line shows upper limit for pSXXF binding affinity (40 µM) due to BRCT variant insolubility (yellow squares) when expressed in *E coli*. Four moderately HSP90-engaged variants are shown as red circles. Known pSXXF disrupting variants at residue R1699 shown as green triangles.

(B) CoIP approach to detect BRCA1-BRCT protein-partner interactions in HEK293T cells using either N-terminally 3x-FLAG tagged full-length BRCA1 (top) or N-terminally 3xFLAG tagged 1,314-BRCA1 truncation (bottom). IP, immunoprecipitation. Neg., negative control.

(C) BRCA1 variant expression using 3xFLAG-BRCA1 cDNA knocked-in at the *HBEGF* locus and dTAG to degrade the endogenous dTAG-BRCA1 in Cal 27 cells. Top panel shows samples treated with 1 µM dTAG-NEG and bottom panel shows 1 µM dTAG indicated by the “+” symbol. BRCA1 antibody targets the N-terminal region of BRCA1. dTAG treatment to degrade dTAG-BRCA1 did not affect 3xFLAG-BRCA1 expression. 3xF, 3xFLAG.

(D) FACS to generate long-term and stable expression of 3xF-BRCA1-2A-mRuby3 mutant cells engineered at the *HBEGF* locus. 3xF-BRCA1-2A-mRuby3 mutant cells were reproducibly dimmer as compared to mRuby3 cells, similar to the wild-type cells.

(E) Maintenance of BRCA1 mutant function for moderately HSP90-engaged variant A1708V was reproduced in two different Cal 27 clones using CellTiter-Glo® to measure cell viability. Relative viability (%) calculated by normalizing dTAG treated cells to matched dTAG-NEG treatments for each cell line separately. Parental cell line is the monoclonally derived homozygous *BRCA1^dTAG/dTAG^* clone.

(F – G) Validation of cell survival assay quantified using crystal violet staining and pixel counting. Shown in (F) is the raw sum density of wells from 96-well plates seeded at low seeding (25 and 100 cells) and high seeding (400 cells) terminated at different days. Horizontal dashed line shows the average value for high cell seeding at days 8 and 9 to illustrate the plateau reached in cell density at high confluency. The linear range of the assay is detectable before the plateau is reached. Shown in (G) is comparisons of relative viability (%) measurements using crystal violet staining or CellTiter-Glo®. Best fit linear regression shown by solid red line and line of identity shown as black dashed line. We consistently observed higher viability measurements using CellTiter-Glo®, presumably because crystal violet processing includes washing steps that remove partially viable cells within the growth medium.

(H – I) PCR-based Sanger sequencing of the BRCT region of 3xFLAG-BRCA1 cDNA variants knocked-in at the *HBEGF* locus. A1708E mutant cells exhibited reduced intensity at the expected band size (arrow) and secondary non-specific bands, suggesting the variant was engineered in the heterozygous state. A1708E mimicked the phenotypic effect of S1715R mutant cells, suggesting the A1708E variant is loss of function consistent with previous reports.^3,5,58,60,69^ Ref., reference BRCA1 sequence.

(J – L) BRCA1 variant-specific function measured by cell survival in HEK293A cells. (L) Cell survival measured using different cell seeding and dTAG treatment conditions to illustrate the reproducibility in partial function for variants moderately engaged by HSP90. Rx 1, 1000 cells seeded and 500 nM dTAG. Rx 2, 1000 cells seed and 1 µM dTAG Day 0 treatment. Rx 3, 1000 cells seeded and 1 µM Day dTAG. Rx 4, 1000 cells seeded and 1 µM dTAG. These HEK293A cells were complemented with a mRuby3 repair template that did not contain an intron carrier, further confirming the robustness of the viability measurements. Statistical significance was determined using Kruskal-Wallis ANOVA test (E and K). \*\*\**p*≤0.001. \*\**p*≤0.01. \*\**p*≤0.05. Data are presented as mean ± standard deviation values from at least two independent experiments or two independent clones.

**Figure S3. HSP90 stabilizes BRCA1 variants in cancer cells, Related to** Figure 3

(A) Cell survival assay in the absence of dTAG treatment to identify non-toxic doses of HSP90 inhibitors for wild-type BRCA1 cells. Each line represents a different cell line with the same dependence for STA-9090 or NVP-HSP990.

(B – C) HSP90-dependent survival of Cal 27 and HEK293A mutant cells expressing moderately (red) or strongly (blue) HSP90-engaged BRCA1 variants. Cells were treated with vehicle DMSO (purple circles), 10 nM STA-9090 (Cal 27) or 2 nM (STA-9090) (red squares). HEK293A cells exhibited higher toxicity to HSP90 inhibitors and was thus treated with lower doses as compared to Cal 27 cells. Null cells were complemented with mRuby3 fluorophore alone. Relative viability (%) quantified by normalizing to dTAG-NEG treated cells for each cell line separately.

(D – F) BRCA1 variant stability in Cal 27 cells depends on HSP90. Cells were treated with 100 µg/mL CHX in the presence of 20 nM STA-9090 (D), DMSO (E), or 39 °C (E) for the indicated number of hours followed by lysis and Western blotting to measure protein turnover. The results showed that the stability of A1708V was more dependent on HSP90 as compared to wild-type BRCA1. D, vehicle (DMSO).

(G) Survival of HEK293A cells expressing BRCA1 variants in the presence of febrile-range heat stress. Cells were grown under basal conditions at 37 °C (purple circles) or at 39 °C (light blue squares).

(H) Heat shock treatment at 39 °C had weak to no effect on Cal 27 and HEK293A mutant cells when dTAG treatment was omitted. Cell survival at 39 °C was normalized to each corresponding mutant cell line at 37 °C, both in the absence of dTAG.

(I) Aggregate detection in mutant cell lines shows equivalent proteostasis effects for cells treated with heat stress at 39 °C. Left panel shows fluorescence intensity for wild-type cells treated with MG132 positive control compound. Right panel shows aggregation propensity factor (APF) calculated by normalizing mean fluorescence intensity between the two conditions indicated by a ‘v’ (vs). FI (AU), fluorescence intensity (arbitrary units).

(J) Cell survival predictions generated by extrapolation / interpolation using HSP90 / HSP70 chaperone interaction magnitude. Our previous report showed that the magnitude of chaperone binding correlates with BRCA1 functional defect (excluding pure contact mutants which were omitted from this analysis).^31^ Large filled squares shows cell survival measurements determine herein (WT, wild-type. AV, A1708V. LR, L1705R. YD, Y1845D. SR, S1715R. AE, A1708E). The large filled circle shows an example variant with the predicted effect colored.

(K) Contour maps illustrating BRCA1-BRCT variant position, HSP90 binding score, and effect on survival at 39 °C. Normalized residue was calculated considering 1,650 and 1,858 as the first and last BRCT domain residues, respectively.

(L – M) Characteristics of HSP90-buffered BRCA1-BRCT variants. Shown in (L) is the sensitivity of mutant cells to HSP90 inhibitor expressed as a difference viability percentage and (M) groups residues exhibiting the strongest response to HSP90 inhibition by secondary structure. The number of variants in each bin is shown in parenthesis above the stacked bar. Statistical significance was determined using multiple two-tailed Mann-Whitney *t*-tests with multiple corrections applied (B, C, G, I and L). \*\*\*\**p*≤0.0001. \*\*\**p*≤0.001. \**p*≤0.05. ns, *p*>0.05. Data are presented as mean ± standard deviation values from at least two independent experiments.

**Figure S4. HSP90-buffered BRCA1 variants exhibit distinct pathogenic properties, Related to** Figure 4

(A) Collected genome instability^58^ assays for BRCA1-BRCT missense variants binned by the magnitude of HSP90 binding. The ‘Other’ bin includes splice site variants,^82^ contact variants,^69^ complex contact / folding variants, and other loss of function variants previously reported.^3,5,31^ Non-buffered variants include those that bind strongly to HSP90 which are expected to be severely misfolded.

(B – C) Cell survival of Cal 27 cancer cells expressing HSP90-buffered BRCA1 variants (red) resist olaparib treatment. Survival was measured by crystal violet staining at 100 nM olaparib. High cell seeding density promotes partial viability of BRCA1-null cells under basal conditions, so that response to olaparib treatment was observable. Higher doses of olaparib sensitized HSP90-buffered variant L1705R, but was moderately toxic to wild-type cells. Ola., olaparib.

(C – D) HSP90_inh_ / PARP_inh_ polytherapy measured by survival assay in Cal 27 cancer cells. Increasing doses of HSP90 inhibitor NVP-HSP990 shown for A1708V and wild-type BRCA1 cells. D, DMSO.

(F – J) Pathogenicity predictions generated by machine learning algorithms.^78–81^ ESM1b predicts were normalized from 0 to 1 using the highest and lowest predicted pathogenicity for BRCA1-BRCT missense variants analyzed. Buffered BRCA1-BRCT variants were defined as moderately-engaged by HSP90 that do not mutate pSXXF coordinating residues. Non-buffered variants comprise mutants strongly-engaged by HSP90 which are prone to severe folding defects. (J) ‘Other’ bind includes pSXXF contact variants, splicing variants, and previously described loss of function variants that do not bind chaperones.^31^ LoF, loss of function. Statistical significance was determined using chi-squared test (A), multiple two-tailed Mann-Whitney *t*-test with multiple corrections applied (B), and Kruskal-Wallis ANOVA test (F–I). \*\*\*\**p*≤0.0001. \*\*\**p*≤0.001. \**p*≤0.01. \**p*≤0.05. ns, no significance. Data are presented as mean ± standard deviation (B) and standard error of the mean (E) values from at least two independent experiments or two independent clones.

## Notes

### Competing Interest Statement

The authors have declared no competing interest.

## REFERENCES

1. Starita, L.M., Young, D.L., Islam, M., Kitzman, J.O., Gullingsrud, J., Hause, R.J., Fowler, D.M., Parvin, J.D., Shendure, J., and Fields, S. (2015). Massively Parallel Functional Analysis of BRCA1 RING Domain Variants. Genetics 200, 413–422. 10.1534/genetics.115.175802.

2. Matreyek, K.A., Starita, L.M., Stephany, J.J., Martin, B., Chiasson, M.A., Gray, V.E., Kircher, M., Khechaduri, A., Dines, J.N., Hause, R.J., et al. (2018). Multiplex assessment of protein variant abundance by massively parallel sequencing. Nat Genet 50, 874–882. 10.1038/s41588-018-0122-z.

3. Findlay, G.M., Daza, R.M., Martin, B., Zhang, M.D., Leith, A.P., Gasperini, M., Janizek, J.D., Huang, X., Starita, L.M., and Shendure, J. (2018). Accurate classification of BRCA1 variants with saturation genome editing. Nature 562, 217–222. 10.1038/s41586-018-0461-z.

4. Fernandes, V.C., Golubeva, V.A., Di Pietro, G., Shields, C., Amankwah, K., Nepomuceno, T.C., de Gregoriis, G., Abreu, R.B.V., Harro, C., Gomes, T.T., et al. (2019). Impact of amino acid substitutions at secondary structures in the BRCT domains of the tumor suppressor BRCA1: Implications for clinical annotation. J Biol Chem 294, 5980–5992. 10.1074/jbc.RA118.005274.

5. Adamovich, A.I., Diabate, M., Banerjee, T., Nagy, G., Smith, N., Duncan, K., Mendoza Mendoza, E., Prida, G., Freitas, M.A., Starita, L.M., and Parvin, J.D. (2022). The functional impact of BRCA1 BRCT domain variants using multiplexed DNA double-strand break repair assays. Am J Hum Genet 109, 618–630. 10.1016/j.ajhg.2022.01.019.

6. Clark, K.A., Paquette, A., Tao, K., Bell, R., Boyle, J.L., Rosenthal, J., Snow, A.K., Stark, A.W., Thompson, B.A., Unger, J., et al. (2022). Comprehensive evaluation and efficient classification of BRCA1 RING domain missense substitutions. Am J Hum Genet 109, 1153–1174. 10.1016/j.ajhg.2022.05.004.

7. Freeman, P.J., Wagstaff, J.F., Fokkema, I.F.A.C., Cutting, G.R., Rehm, H.L., Davies, A.C., den Dunnen, J.T., Gretton, L.J., and Dalgleish, R. (2024). Standardizing variant naming in literature with VariantValidator to increase diagnostic rates. Nat Genet. 10.1038/s41588-024-01938-w.

8. López-Otín, C., Blasco, M.A., Partridge, L., Serrano, M., and Kroemer, G. (2013). The hallmarks of aging. Cell 153, 1194–1217. 10.1016/j.cell.2013.05.039.

9. Mo, C., Hannan, A.J., and Renoir, T. (2015). Environmental factors as modulators of neurodegeneration: insights from gene-environment interactions in Huntington’s disease. Neurosci Biobehav Rev 52, 178–192. 10.1016/j.neubiorev.2015.03.003.

10. Tan, S.L.W., Chadha, S., Liu, Y., Gabasova, E., Perera, D., Ahmed, K., Constantinou, S., Renaudin, X., Lee, M., Aebersold, R., and Venkitaraman, A.R. (2017). A Class of Environmental and Endogenous Toxins Induces BRCA2 Haploinsufficiency and Genome Instability. Cell 169, 1105–1118.e1115. 10.1016/j.cell.2017.05.010.

11. Hingorani, A.D., Gratton, J., Finan, C., Schmidt, A.F., Patel, R., Sofat, R., Kuan, V., Langenberg, C., Hemingway, H., Morris, J.K., and Wald, N.J. (2023). Performance of polygenic risk scores in screening, prediction, and risk stratification: secondary analysis of data in the Polygenic Score Catalog. BMJ Med 2, e000554. 10.1136/bmjmed-2023-000554.

12. Burke, W., Aitken, M.L., Chen, S.H., and Scott, C.R. (1992). Variable severity of pulmonary disease in adults with identical cystic fibrosis mutations. Chest 102, 506–509. 10.1378/chest.102.2.506.

13. Prevalence and penetrance of BRCA1 and BRCA2 mutations in a population-based series of breast cancer cases. Anglian Breast Cancer Study Group. (2000). Br J Cancer 83, 1301–1308. 10.1054/bjoc.2000.1407.

14. Weatherall, D.J. (2001). Phenotype-genotype relationships in monogenic disease: lessons from the thalassaemias. Nat Rev Genet 2, 245–255. 10.1038/35066048.

15. Schreiner, S., Cossais, F., Fischer, K., Scholz, S., Bösl, M.R., Holtmann, B., Sendtner, M., and Wegner, M. (2007). Hypomorphic Sox10 alleles reveal novel protein functions and unravel developmental differences in glial lineages. Development 134, 3271–3281. 10.1242/dev.003350.

16. Perlis, R.H., Smoller, J.W., Mysore, J., Sun, M., Gillis, T., Purcell, S., Rietschel, M., Nöthen, M.M., Witt, S., Maier, W., et al. (2010). Prevalence of incompletely penetrant Huntington’s disease alleles among individuals with major depressive disorder. Am J Psychiatry 167, 574–579. 10.1176/appi.ajp.2009.09070973.

17. Avila, E.M., Uzel, G., Hsu, A., Milner, J.D., Turner, M.L., Pittaluga, S., Freeman, A.F., and Holland, S.M. (2010). Highly variable clinical phenotypes of hypomorphic RAG1 mutations. Pediatrics 126, e1248–1252. 10.1542/peds.2009-3171.

18. Cooper, D.N., Krawczak, M., Polychronakos, C., Tyler-Smith, C., and Kehrer-Sawatzki, H. (2013). Where genotype is not predictive of phenotype: towards an understanding of the molecular basis of reduced penetrance in human inherited disease. Hum Genet 132, 1077–1130. 10.1007/s00439-013-1331-2.

19. Meshaal, S.S., El Hawary, R.E., Abd Elaziz, D.S., Eldash, A., Alkady, R., Lotfy, S., Mauracher, A.A., Opitz, L., Pachlopnik Schmid, J., van der Burg, M., et al. (2019). Phenotypical heterogeneity in RAG-deficient patients from a highly consanguineous population. Clin Exp Immunol 195, 202–212. 10.1111/cei.13222.

20. Pei, J., Kinch, L.N., Otwinowski, Z., and Grishin, N.V. (2020). Mutation severity spectrum of rare alleles in the human genome is predictive of disease type. PLoS Comput Biol 16, e1007775. 10.1371/journal.pcbi.1007775.

21. Gudmundsson, S., Singer-Berk, M., Stenton, S.L., Goodrich, J.K., Wilson, M.W., Einson, J., Watts, N.A., Lappalainen, T., Rehm, H.L., MacArthur, D.G., et al. (2024). Exploring penetrance of clinically relevant variants in over 800,000 humans from the Genome Aggregation Database. bioRxiv. 10.1101/2024.06.12.593113.

22. Dill, K.A., and Chan, H.S. (1997). From Levinthal to pathways to funnels. Nat Struct Biol 4, 10–19.

23. Fersht, A. (1998). Structure and Mechanism in Protein Science: A Guide to Enzyme Catalysis and Protein Folding (W.H. Freeman).

24. Taverna, D.M., and Goldstein, R.A. (2002). Why are proteins marginally stable? Proteins 46, 105–109. 10.1002/prot.10016.

25. Hamborg, L., Horsted, E.W., Johansson, K.E., Willemoës, M., Lindorff-Larsen, K., and Teilum, K. (2020). Global analysis of protein stability by temperature and chemical denaturation. Anal Biochem 605, 113863. 10.1016/j.ab.2020.113863.

26. Dutta, P., Roy, P., and Sengupta, N. (2022). Effects of External Perturbations on Protein Systems: A Microscopic View. ACS Omega 7, 44556–44572. 10.1021/acsomega.2c06199.

27. Ó’Fágáin, C. (2023). Protein Stability: Enhancement and Measurement. Methods Mol Biol 2699, 369–419. 10.1007/978-1-0716-3362-5_18.

28. Sahni, N., Yi, S., Taipale, M., Fuxman Bass, J.I., Coulombe-Huntington, J., Yang, F., Peng, J., Weile, J., Karras, G.I., Wang, Y., et al. (2015). Widespread macromolecular interaction perturbations in human genetic disorders. Cell 161, 647–660. 10.1016/j.cell.2015.04.013.

29. Lacoste, J., Haghighi, M., Haider, S., Lin, Z.Y., Segal, D., Reno, C., Qian, W.W., Xiong, X., Shafqat-Abbasi, H., Ryder, P.V., et al. (2023). Pervasive mislocalization of pathogenic coding variants underlying human disorders. bioRxiv. 10.1101/2023.09.05.556368.

30. Weng, C., Faure, A.J., Escobedo, A., and Lehner, B. (2023). The energetic and allosteric landscape for KRAS inhibition. Nature. 10.1038/s41586-023-06954-0.

31. Gracia, B., Montes, P., Gutierrez, A.M., Arun, B., and Karras, G.I. (2024). Protein-folding chaperones predict structure-function relationships and cancer risk in BRCA1 mutation carriers. Cell Rep 43, 113803. 10.1016/j.celrep.2024.113803.

32. Bloom, J.D., Labthavikul, S.T., Otey, C.R., and Arnold, F.H. (2006). Protein stability promotes evolvability. Proc Natl Acad Sci U S A 103, 5869–5874. 10.1073/pnas.0510098103.

33. Zabinsky, R.A., Mason, G.A., Queitsch, C., and Jarosz, D.F. (2018). It’s not magic - Hsp90 and its effects on genetic and epigenetic variation. Semin Cell Dev Biol. 10.1016/j.semcdb.2018.05.015.

34. Borkovich, K.A., Farrelly, F.W., Finkelstein, D.B., Taulien, J., and Lindquist, S. (1989). hsp82 is an essential protein that is required in higher concentrations for growth of cells at higher temperatures. Mol Cell Biol 9, 3919–3930.

35. Bhattacharya, K., Maiti, S., Zahoran, S., Weidenauer, L., Hany, D., Wider, D., Bernasconi, L., Quadroni, M., Collart, M., and Picard, D. (2022). Translational reprogramming in response to accumulating stressors ensures critical threshold levels of Hsp90 for mammalian life. Nat Commun 13, 6271. 10.1038/s41467-022-33916-3.

36. Kikis, E.A., Gidalevitz, T., and Morimoto, R.I. (2010). Protein homeostasis in models of aging and age-related conformational disease. Adv Exp Med Biol 694, 138–159. 10.1007/978-1-4419-7002-2_11.

37. Hartl, F.U., Bracher, A., and Hayer-Hartl, M. (2011). Molecular chaperones in protein folding and proteostasis. Nature 475, 324–332. 10.1038/nature10317.

38. Guo, J., Huang, X., Dou, L., Yan, M., Shen, T., Tang, W., and Li, J. (2022). Aging and aging-related diseases: from molecular mechanisms to interventions and treatments. Signal Transduct Target Ther 7, 391. 10.1038/s41392-022-01251-0.

39. Rutherford, S.L., and Lindquist, S. (1998). Hsp90 as a capacitor for morphological evolution. Nature 396, 336–342. 10.1038/24550.

40. Queitsch, C., Sangster, T.A., and Lindquist, S. (2002). Hsp90 as a capacitor of phenotypic variation. Nature 417, 618–624. 10.1038/nature749.

41. Cowen, L.E., Singh, S.D., Köhler, J.R., Collins, C., Zaas, A.K., Schell, W.A., Aziz, H., Mylonakis, E., Perfect, J.R., Whitesell, L., and Lindquist, S. (2009). Harnessing Hsp90 function as a powerful, broadly effective therapeutic strategy for fungal infectious disease. Proc Natl Acad Sci U S A 106, 2818–2823. 10.1073/pnas.0813394106.

42. Jarosz, D.F., and Lindquist, S. (2010). Hsp90 and environmental stress transform the adaptive value of natural genetic variation. Science 330, 1820–1824. 10.1126/science.1195487.

43. Rohner, N., Jarosz, D.F., Kowalko, J.E., Yoshizawa, M., Jeffery, W.R., Borowsky, R.L., Lindquist, S., and Tabin, C.J. (2013). Cryptic variation in morphological evolution: HSP90 as a capacitor for loss of eyes in cavefish. Science 342, 1372–1375. 10.1126/science.1240276.

44. Revie, N.M., Iyer, K.R., Robbins, N., and Cowen, L.E. (2018). Antifungal drug resistance: evolution, mechanisms and impact. Curr Opin Microbiol 45, 70–76. 10.1016/j.mib.2018.02.005.

45. Condic, N., Amiji, H., Patel, D., Shropshire, W.C., Lermi, N.O., Sabha, Y., John, B., Hanson, B., and Karras, G.I. (2024). Selection for robust metabolism in domesticated yeasts is driven by adaptation to Hsp90 stress. Science 385, eadi3048. 10.1126/science.adi3048.

46. Karras, G.I., Yi, S., Sahni, N., Fischer, M., Xie, J., Vidal, M., D’Andrea, A.D., Whitesell, L., and Lindquist, S. (2017). HSP90 Shapes the Consequences of Human Genetic Variation. Cell 168, 856–866.e812. 10.1016/j.cell.2017.01.023.

47. Miki, Y., Swensen, J., Shattuck-Eidens, D., Futreal, P.A., Harshman, K., Tavtigian, S., Liu, Q., Cochran, C., Bennett, L.M., and Ding, W. (1994). A strong candidate for the breast and ovarian cancer susceptibility gene BRCA1. Science 266, 66–71. 10.1126/science.7545954.

48. Friedman, L.S., Ostermeyer, E.A., Szabo, C.I., Dowd, P., Lynch, E.D., Rowell, S.E., and King, M.C. (1994). Confirmation of BRCA1 by analysis of germline mutations linked to breast and ovarian cancer in ten families. Nat Genet 8, 399–404. 10.1038/ng1294-399.

49. Ashworth, A. (2024). Thirty years since the race to the BRCA1 gene. Nature 634, 1062–1063. 10.1038/d41586-024-03358-6.

50. Drost, R., Bouwman, P., Rottenberg, S., Boon, U., Schut, E., Klarenbeek, S., Klijn, C., van der Heijden, I., van der Gulden, H., Wientjens, E., et al. (2011). BRCA1 RING function is essential for tumor suppression but dispensable for therapy resistance. Cancer Cell 20, 797–809. 10.1016/j.ccr.2011.11.014.

51. Liu, Y., and Lu, L.Y. (2020). BRCA1 and homologous recombination: implications from mouse embryonic development. Cell Biosci 10, 49. 10.1186/s13578-020-00412-4.

52. Leal, A.F., Herreno-Pachón, A.M., Benincore-Flórez, E., Karunathilaka, A., and Tomatsu, S. (2024). Current Strategies for Increasing Knock-In Efficiency in CRISPR/Cas9-Based Approaches. Int J Mol Sci 25. 10.3390/ijms25052456.

53. Nabet, B., Roberts, J.M., Buckley, D.L., Paulk, J., Dastjerdi, S., Yang, A., Leggett, A.L., Erb, M.A., Lawlor, M.A., Souza, A., et al. (2018). The dTAG system for immediate and target-specific protein degradation. Nat Chem Biol 14, 431–441. 10.1038/s41589-018-0021-8.

54. Nabet, B., Ferguson, F.M., Seong, B.K.A., Kuljanin, M., Leggett, A.L., Mohardt, M.L., Robichaud, A., Conway, A.S., Buckley, D.L., Mancias, J.D., et al. (2020). Rapid and direct control of target protein levels with VHL-recruiting dTAG molecules. Nat Commun 11, 4687. 10.1038/s41467-020-18377-w.

55. Oliveira-Costa, J.P., Oliveira, L.R., Zanetti, R., Zanetti, J.S., da Silveira, G.G., Chavichiolli Buim, M.E., Zucoloto, S., Ribeiro-Silva, A., and Soares, F.A. (2014). BRCA1 and γH2AX as independent prognostic markers in oral squamous cell carcinoma. Oncoscience 1, 383–391. 10.18632/oncoscience.47.

56. Payungwong, T., Angkulkrerkkrai, K., Chaiboonchoe, A., Lausoontornsiri, W., Jirawatnotai, S., and Chindavijak, S. (2024). Comparison of mutation landscapes of pretreatment versus recurrent squamous cell carcinoma of the oral cavity: The possible mechanism of resistance to standard treatment. Cancer Rep (Hoboken) 7, e2004. 10.1002/cnr2.2004.

57. Li, S., Akrap, N., Cerboni, S., Porritt, M.J., Wimberger, S., Lundin, A., Möller, C., Firth, M., Gordon, E., Lazovic, B., et al. (2021). Universal toxin-based selection for precise genome engineering in human cells. Nat Commun 12, 497. 10.1038/s41467-020-20810-z.

58. Cusin, I., Teixeira, D., Zahn-Zabal, M., Rech de Laval, V., Gleizes, A., Viassolo, V., Chappuis, P.O., Hutter, P., Bairoch, A., and Gaudet, P. (2018). A new bioinformatics tool to help assess the significance of BRCA1 variants. Hum Genomics 12, 36. 10.1186/s40246-018-0168-0.

59. Wu, Q., Jubb, H., and Blundell, T.L. (2015). Phosphopeptide interactions with BRCA1 BRCT domains: More than just a motif. Prog Biophys Mol Biol 117, 143–148. 10.1016/j.pbiomolbio.2015.02.003.

60. Williams, R.S., Chasman, D.I., Hau, D.D., Hui, B., Lau, A.Y., and Glover, J.N. (2003). Detection of protein folding defects caused by BRCA1-BRCT truncation and missense mutations. J Biol Chem 278, 53007–53016. 10.1074/jbc.M310182200.

61. Lee, M.S., Green, R., Marsillac, S.M., Coquelle, N., Williams, R.S., Yeung, T., Foo, D., Hau, D.D., Hui, B., Monteiro, A.N., and Glover, J.N. (2010). Comprehensive analysis of missense variations in the BRCT domain of BRCA1 by structural and functional assays. Cancer Res 70, 4880–4890. 10.1158/0008-5472.CAN-09-4563.

62. Rowling, P.J., Cook, R., and Itzhaki, L.S. (2010). Toward classification of BRCA1 missense variants using a biophysical approach. J Biol Chem 285, 20080–20087. 10.1074/jbc.M109.088922.

63. Landrum, M.J., Lee, J.M., Benson, M., Brown, G.R., Chao, C., Chitipiralla, S., Gu, B., Hart, J., Hoffman, D., Jang, W., et al. (2018). ClinVar: improving access to variant interpretations and supporting evidence. Nucleic Acids Res 46, D1062–D1067. 10.1093/nar/gkx1153.

64. Yu, X., Chini, C.C., He, M., Mer, G., and Chen, J. (2003). The BRCT domain is a phospho-protein binding domain. Science 302, 639–642. 10.1126/science.1088753.

65. Coquelle, N., Green, R., and Glover, J.N. (2011). Impact of BRCA1 BRCT domain missense substitutions on phosphopeptide recognition. Biochemistry 50, 4579–4589. 10.1021/bi2003795.

66. Nair, S.C., Rimerman, R.A., Toran, E.J., Chen, S., Prapapanich, V., Butts, R.N., and Smith, D.F. (1997). Molecular cloning of human FKBP51 and comparisons of immunophilin interactions with Hsp90 and progesterone receptor. Mol Cell Biol 17, 594–603. 10.1128/MCB.17.2.594.

67. Li, S., Chen, P.L., Subramanian, T., Chinnadurai, G., Tomlinson, G., Osborne, C.K., Sharp, Z.D., and Lee, W.H. (1999). Binding of CtIP to the BRCT repeats of BRCA1 involved in the transcription regulation of p21 is disrupted upon DNA damage. J Biol Chem 274, 11334–11338. 10.1074/jbc.274.16.11334.

68. Wu, Q., Paul, A., Su, D., Mehmood, S., Foo, T.K., Ochi, T., Bunting, E.L., Xia, B., Robinson, C.V., Wang, B., and Blundell, T.L. (2016). Structure of BRCA1-BRCT/Abraxas Complex Reveals Phosphorylation-Dependent BRCT Dimerization at DNA Damage Sites. Mol Cell 61, 434–448. 10.1016/j.molcel.2015.12.017.

69. Jhuraney, A., Velkova, A., Johnson, R.C., Kessing, B., Carvalho, R.S., Whiley, P., Spurdle, A.B., Vreeswijk, M.P., Caputo, S.M., Millot, G.A., et al. (2015). BRCA1 Circos: a visualisation resource for functional analysis of missense variants. J Med Genet 52, 224–230. 10.1136/jmedgenet-2014-102766.

70. Li, Z.N., and Luo, Y. (2023). HSP90 inhibitors and cancer: Prospects for use in targeted therapies (Review). Oncol Rep 49. 10.3892/or.2022.8443.

71. Stecklein, S.R., Kumaraswamy, E., Behbod, F., Wang, W., Chaguturu, V., Harlan-Williams, L.M., and Jensen, R.A. (2012). BRCA1 and HSP90 cooperate in homologous and non-homologous DNA double-strand-break repair and G2/M checkpoint activation. Proc Natl Acad Sci U S A 109, 13650–13655. 10.1073/pnas.1203326109.

72. 72. Lesire, L., Chaput, L., Cruz De Casas, P., Rousseau, F., Piveteau, C., Dumont, J., Pointu, D., Déprez, B., and Leroux, F. (2020). High-Throughput Image-Based Aggresome Quantification. SLAS Discov 25, 783–791. 10.1177/2472555220919708.

73. Zhao, P., Wang, C., Sun, S., Wang, X., and Balch, W.E. (2024). Tracing genetic diversity captures the molecular basis of misfolding disease. Nat Commun 15, 3333. 10.1038/s41467- 024-47520-0.

74. Farmer, H., McCabe, N., Lord, C.J., Tutt, A.N., Johnson, D.A., Richardson, T.B., Santarosa, M., Dillon, K.J., Hickson, I., Knights, C., et al. (2005). Targeting the DNA repair defect in BRCA mutant cells as a therapeutic strategy. Nature 434, 917–921. 10.1038/nature03445.

75. Johnson, N., Johnson, S.F., Yao, W., Li, Y.C., Choi, Y.E., Bernhardy, A.J., Wang, Y., Capelletti, M., Sarosiek, K.A., Moreau, L.A., et al. (2013). Stabilization of mutant BRCA1 protein confers PARP inhibitor and platinum resistance. Proc Natl Acad Sci U S A 110, 17041–17046. 10.1073/pnas.1305170110.

76. Ragupathi, A., Singh, M., Perez, A.M., and Zhang, D. (2023). Targeting the BRCA1/2 deficient cancer with PARP inhibitors: Clinical outcomes and mechanistic insights. Front Cell Dev Biol 11, 1133472. 10.3389/fcell.2023.1133472.

77. Nepomuceno, T.C., Foo, T.K., Richardson, M.E., Ranola, J.M.O., Weyandt, J., Varga, M.J., Alarcon, A., Gutierrez, D., von Wachenfeldt, A., Eriksson, D., et al. (2023). BRCA1 frameshift variants leading to extended incorrect protein C termini. HGG Adv 4, 100240. 10.1016/j.xhgg.2023.100240.

78. Frazer, J., Notin, P., Dias, M., Gomez, A., Min, J.K., Brock, K., Gal, Y., and Marks, D.S. (2021). Disease variant prediction with deep generative models of evolutionary data. Nature 599, 91–95. 10.1038/s41586-021-04043-8.

79. Wu, Y., Liu, H., Li, R., Sun, S., Weile, J., and Roth, F.P. (2021). Improved pathogenicity prediction for rare human missense variants. Am J Hum Genet 108, 2389. 10.1016/j.ajhg.2021.11.010.

80. Cheng, J., Novati, G., Pan, J., Bycroft, C., Žemgulytė, A., Applebaum, T., Pritzel, A., Wong, L.H., Zielinski, M., Sargeant, T., et al. (2023). Accurate proteome-wide missense variant effect prediction with AlphaMissense. Science 381, eadg7492. 10.1126/science.adg7492.

81. Brandes, N., Goldman, G., Wang, C.H., Ye, C.J., and Ntranos, V. (2023). Genome-wide prediction of disease variant effects with a deep protein language model. Nat Genet 55, 1512–1522. 10.1038/s41588-023-01465-0.

82. Ahlborn, L.B., Dandanell, M., Steffensen, A.Y., Jønson, L., Nielsen, F.C., and Hansen, T.V. (2015). Splicing analysis of 14 BRCA1 missense variants classifies nine variants as pathogenic. Breast Cancer Res Treat 150, 289–298. 10.1007/s10549-015-3313-7.

83. Cerami, E., Gao, J., Dogrusoz, U., Gross, B.E., Sumer, S.O., Aksoy, B.A., Jacobsen, A., Byrne, C.J., Heuer, M.L., Larsson, E., et al. (2012). The cBio cancer genomics portal: an open platform for exploring multidimensional cancer genomics data. Cancer Discov 2, 401–404. 10.1158/2159-8290.CD-12-0095.

84. Gao, J., Aksoy, B.A., Dogrusoz, U., Dresdner, G., Gross, B., Sumer, S.O., Sun, Y., Jacobsen, A., Sinha, R., Larsson, E., et al. (2013). Integrative analysis of complex cancer genomics and clinical profiles using the cBioPortal. Sci Signal 6, pl1. 10.1126/scisignal.2004088.

85. Karczewski, K.J., Francioli, L.C., Tiao, G., Cummings, B.B., Alföldi, J., Wang, Q., Collins, R.L., Laricchia, K.M., Ganna, A., Birnbaum, D.P., et al. (2020). The mutational constraint spectrum quantified from variation in 141,456 humans. Nature 581, 434–443. 10.1038/s41586-020-2308-7.

86. Fokkema, I.F., Taschner, P.E., Schaafsma, G.C., Celli, J., Laros, J.F., and den Dunnen, J.T. (2011). LOVD v.2.0: the next generation in gene variant databases. Hum Mutat 32, 557–563. 10.1002/humu.21438.

87. Cowen, L.E., and Lindquist, S. (2005). Hsp90 potentiates the rapid evolution of new traits: drug resistance in diverse fungi. Science 309, 2185–2189. 10.1126/science.1118370.

88. Sangster, T.A., Salathia, N., Lee, H.N., Watanabe, E., Schellenberg, K., Morneau, K., Wang, H., Undurraga, S., Queitsch, C., and Lindquist, S. (2008). HSP90-buffered genetic variation is common in Arabidopsis thaliana. Proc Natl Acad Sci U S A 105, 2969–2974. 10.1073/pnas.0712210105.

89. Chen, S., and Parmigiani, G. (2007). Meta-analysis of BRCA1 and BRCA2 penetrance. J Clin Oncol 25, 1329–1333. 10.1200/JCO.2006.09.1066.

90. van der Kolk, D.M., de Bock, G.H., Leegte, B.K., Schaapveld, M., Mourits, M.J., de Vries, J., van der Hout, A.H., and Oosterwijk, J.C. (2010). Penetrance of breast cancer, ovarian cancer and contralateral breast cancer in BRCA1 and BRCA2 families: high cancer incidence at older age. Breast Cancer Res Treat 124, 643–651. 10.1007/s10549-010-0805-3.

91. Keung, M.Y., Wu, Y., Badar, F., and Vadgama, J.V. (2020). Response of Breast Cancer Cells to PARP Inhibitors Is Independent of BRCA Status. J Clin Med 9. 10.3390/jcm9040940.

92. Tobalina, L., Armenia, J., Irving, E., O’Connor, M.J., and Forment, J.V. (2021). A meta-analysis of reversion mutations in BRCA genes identifies signatures of DNA end-joining repair mechanisms driving therapy resistance. Ann Oncol 32, 103–112. 10.1016/j.annonc.2020.10.470.

93. Makhnoon, S., Chen, M., Levin, B., Ensinger, M., Mattie, K.D., Grana, G., Shete, S., Arun, B.K., and Peterson, S.K. (2022). Use of breast surveillance between women with pathogenic variants and variants of uncertain significance in breast cancer susceptibility genes. Cancer 128, 3709–3717. 10.1002/cncr.34429.

94. Kingdom, R., and Wright, C.F. (2022). Incomplete Penetrance and Variable Expressivity: From Clinical Studies to Population Cohorts. Front Genet 13, 920390. 10.3389/fgene.2022.920390.

95. Jackson, L.M., and Moldovan, G.L. (2022). Mechanisms of PARP1 inhibitor resistance and their implications for cancer treatment. NAR Cancer 4, zcac042. 10.1093/narcan/zcac042.

96. Murciano-Goroff, Y.R., Schram, A.M., Rosen, E.Y., Won, H., Gong, Y., Noronha, A.M., Janjigian, Y.Y., Stadler, Z.K., Chang, J.C., Yang, S.R., et al. (2022). Reversion mutations in germline BRCA1/2-mutant tumors reveal a BRCA-mediated phenotype in non-canonical histologies. Nat Commun 13, 7182. 10.1038/s41467-022-34109-8.

97. Noddings, C.M., Wang, R.Y., Johnson, J.L., and Agard, D.A. (2022). Structure of Hsp90-p23- GR reveals the Hsp90 client-remodelling mechanism. Nature 601, 465–469. 10.1038/s41586-021-04236-1.

98. Wang, R.Y., Noddings, C.M., Kirschke, E., Myasnikov, A.G., Johnson, J.L., and Agard, D.A. (2022). Structure of Hsp90-Hsp70-Hop-GR reveals the Hsp90 client-loading mechanism. Nature 601, 460–464. 10.1038/s41586-021-04252-1.

99. Lovelock, P.K., Spurdle, A.B., Mok, M.T., Farrugia, D.J., Lakhani, S.R., Healey, S., Arnold, S., Buchanan, D., Couch, F.J., Henderson, B.R., et al. (2007). Identification of BRCA1 missense substitutions that confer partial functional activity: potential moderate risk variants? Breast Cancer Res 9, R82. 10.1186/bcr1826.

100. Spurdle, A.B., Whiley, P.J., Thompson, B., Feng, B., Healey, S., Brown, M.A., Pettigrew, C., Van Asperen, C.J., Ausems, M.G., Kattentidt-Mouravieva, A.A., et al. (2012). BRCA1 R1699Q variant displaying ambiguous functional abrogation confers intermediate breast and ovarian cancer risk. J Med Genet 49, 525–532. 10.1136/jmedgenet-2012-101037.

101. Moghadasi, S., Meeks, H.D., Vreeswijk, M.P., Janssen, L.A., Borg, Å., Ehrencrona, H., Paulsson-Karlsson, Y., Wappenschmidt, B., Engel, C., Gehrig, A., et al. (2018). The BRCA1 c. 5096G>A p.Arg1699Gln (R1699Q) intermediate risk variant: breast and ovarian cancer risk estimation and recommendations for clinical management from the ENIGMA consortium. J Med Genet 55, 15–20. 10.1136/jmedgenet-2017-104560.

102. Carvalho, M.A., Billack, B., Chan, E., Worley, T., Cayanan, C., and Monteiro, A.N. (2002). Mutations in the BRCT domain confer temperature sensitivity to BRCA1 in transcription activation. Cancer Biol Ther 1, 502–508. 10.4161/cbt.1.5.165.

103. Worley, T., Vallon-Christersson, J., Billack, B., Borg, A., and Monteiro, A.N. (2002). A naturally occurring allele of BRCA1 coding for a temperature-sensitive mutant protein. Cancer Biol Ther 1, 497–501. 10.4161/cbt.1.5.164.

104. Pillai, R.N., and Ramalingam, S.S. (2018). Throwing More Cold Water on Heat Shock Protein 90 Inhibitors in NSCLC. J Thorac Oncol 13, 473–474. 10.1016/j.jtho.2018.02.010.

105. Gao, C., Peng, Y.N., Wang, H.Z., Fang, S.L., Zhang, M., Zhao, Q., and Liu, J. (2019). Inhibition of Heat Shock Protein 90 as a Novel Platform for the Treatment of Cancer. Curr Pharm Des 25, 849–855. 10.2174/1381612825666190503145944.

106. Lang, J.E., Forero-Torres, A., Yee, D., Yau, C., Wolf, D., Park, J., Parker, B.A., Chien, A.J., Wallace, A.M., Murthy, R., et al. (2022). Safety and efficacy of HSP90 inhibitor ganetespib for neoadjuvant treatment of stage II/III breast cancer. NPJ Breast Cancer 8, 128. 10.1038/s41523-022-00493-z.

107. Rastogi, S., Joshi, A., Sato, N., Lee, S., Lee, M.J., Trepel, J.B., and Neckers, L. (2024). An update on the status of HSP90 inhibitors in cancer clinical trials. Cell Stress Chaperones 29, 519–539. 10.1016/j.cstres.2024.05.005.

108. Li, H., Liu, Z.Y., Wu, N., Chen, Y.C., Cheng, Q., and Wang, J. (2020). PARP inhibitor resistance: the underlying mechanisms and clinical implications. Mol Cancer 19, 107. 10.1186/s12943-020-01227-0.

109. Dias, M.P., Moser, S.C., Ganesan, S., and Jonkers, J. (2021). Understanding and overcoming resistance to PARP inhibitors in cancer therapy. Nat Rev Clin Oncol 18, 773–791. 10.1038/s41571-021-00532-x.

110. Giudice, E., Gentile, M., Salutari, V., Ricci, C., Musacchio, L., Carbone, M.V., Ghizzoni, V., Camarda, F., Tronconi, F., Nero, C., et al. (2022). PARP Inhibitors Resistance: Mechanisms and Perspectives. Cancers (Basel) 14. 10.3390/cancers14061420.

111. Ewers, K.M., Patil, S., Kopp, W., Thomale, J., Quilitz, T., Magerhans, A., Wang, X., Hessmann, E., and Dobbelstein, M. (2021). HSP90 Inhibition Synergizes with Cisplatin to Eliminate Basal-like Pancreatic Ductal Adenocarcinoma Cells. Cancers (Basel) 13. 10.3390/cancers13246163.

112. Liu, L., Deng, Y., Zheng, Z., Deng, Z., Zhang, J., Li, J., Liang, M., Zhou, X., Tan, W., Yang, H., et al. (2021). Hsp90 Inhibitor STA9090 Sensitizes Hepatocellular Carcinoma to Hyperthermia-Induced DNA Damage by Suppressing DNA-PKcs Protein Stability and mRNA Transcription. Mol Cancer Ther 20, 1880–1892. 10.1158/1535-7163.MCT-21-0215.

113. Schymkowitz, J., Borg, J., Stricher, F., Nys, R., Rousseau, F., and Serrano, L. (2005). The FoldX web server: an online force field. Nucleic Acids Res 33, W382–388. 10.1093/nar/gki387.

114. Shiozaki, E.N., Gu, L., Yan, N., and Shi, Y. (2004). Structure of the BRCT repeats of BRCA1 bound to a BACH1 phosphopeptide: implications for signaling. Mol Cell 14, 405–412. 10.1016/s1097-2765(04)00238-2.

115. Atchley, D.P., Albarracin, C.T., Lopez, A., Valero, V., Amos, C.I., Gonzalez-Angulo, A.M., Hortobagyi, G.N., and Arun, B.K. (2008). Clinical and pathologic characteristics of patients with BRCA-positive and BRCA-negative breast cancer. J Clin Oncol 26, 4282–4288. 10.1200/JCO.2008.16.6231.

116. Sanford, R.A., Song, J., Gutierrez-Barrera, A.M., Profato, J., Woodson, A., Litton, J.K., Bedrosian, I., Albarracin, C.T., Valero, V., and Arun, B. (2015). High incidence of germline BRCA mutation in patients with ER low-positive/PR low-positive/HER-2 neu negative tumors. Cancer 121, 3422–3427. 10.1002/cncr.29572.

117. Crowley, L.C., Christensen, M.E., and Waterhouse, N.J. (2016). Measuring Survival of Adherent Cells with the Colony-Forming Assay. Cold Spring Harb Protoc 2016. 10.1101/pdb.prot087171.

118. Williams, R.S., Lee, M.S., Hau, D.D., and Glover, J.N. (2004). Structural basis of phosphopeptide recognition by the BRCT domain of BRCA1. Nat Struct Mol Biol 11, 519–525. 10.1038/nsmb776.

119. Hayryan, S., Hu, C.K., Skrivánek, J., Hayryane, E., and Pokorný, I. (2005). A new analytical method for computing solvent-accessible surface area of macromolecules and its gradients. J Comput Chem 26, 334–343. 10.1002/jcc.20125.

120. Heinig, M., and Frishman, D. (2004). STRIDE: a web server for secondary structure assignment from known atomic coordinates of proteins. Nucleic Acids Res 32, W500–502. 10.1093/nar/gkh429.

121. Ferruz, N., Schmidt, S., and Höcker, B. (2021). ProteinTools: a toolkit to analyze protein structures. Nucleic Acids Res 49, W559–W566. 10.1093/nar/gkab375.

