## Supplementary material for "HSP90 buffers deleterious genetic variations in *BRCA1*": Figure S1

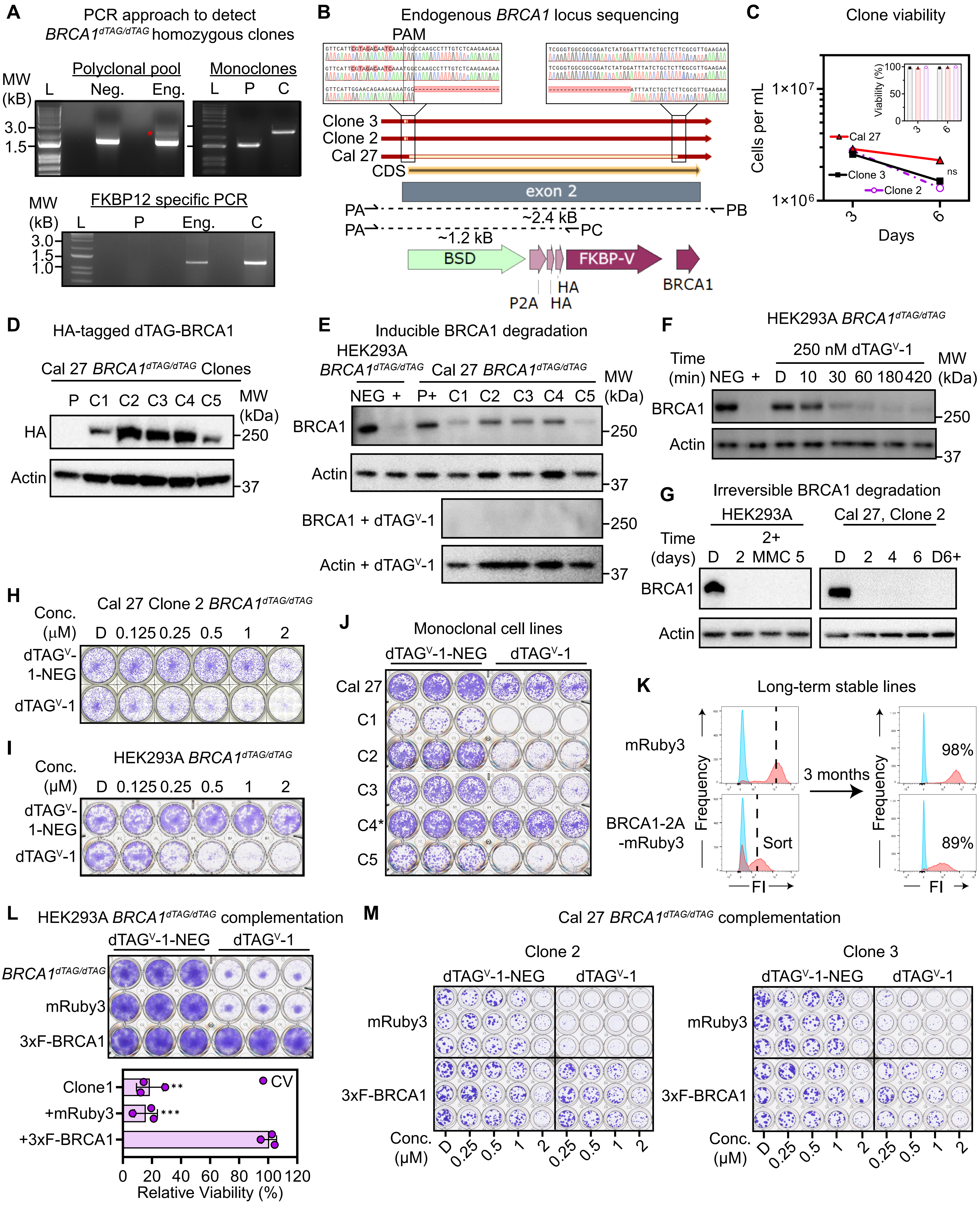


**Figure S1. $BRCA1^{dTAG/dTAG}$ knock-ins enable conditional complementation of *BRCA1* in multiple cell lines, Related to Figure 1**
