## Supplementary figures and images for "HSP90 buffers deleterious genetic variations in *BRCA1*"

### Figure S2

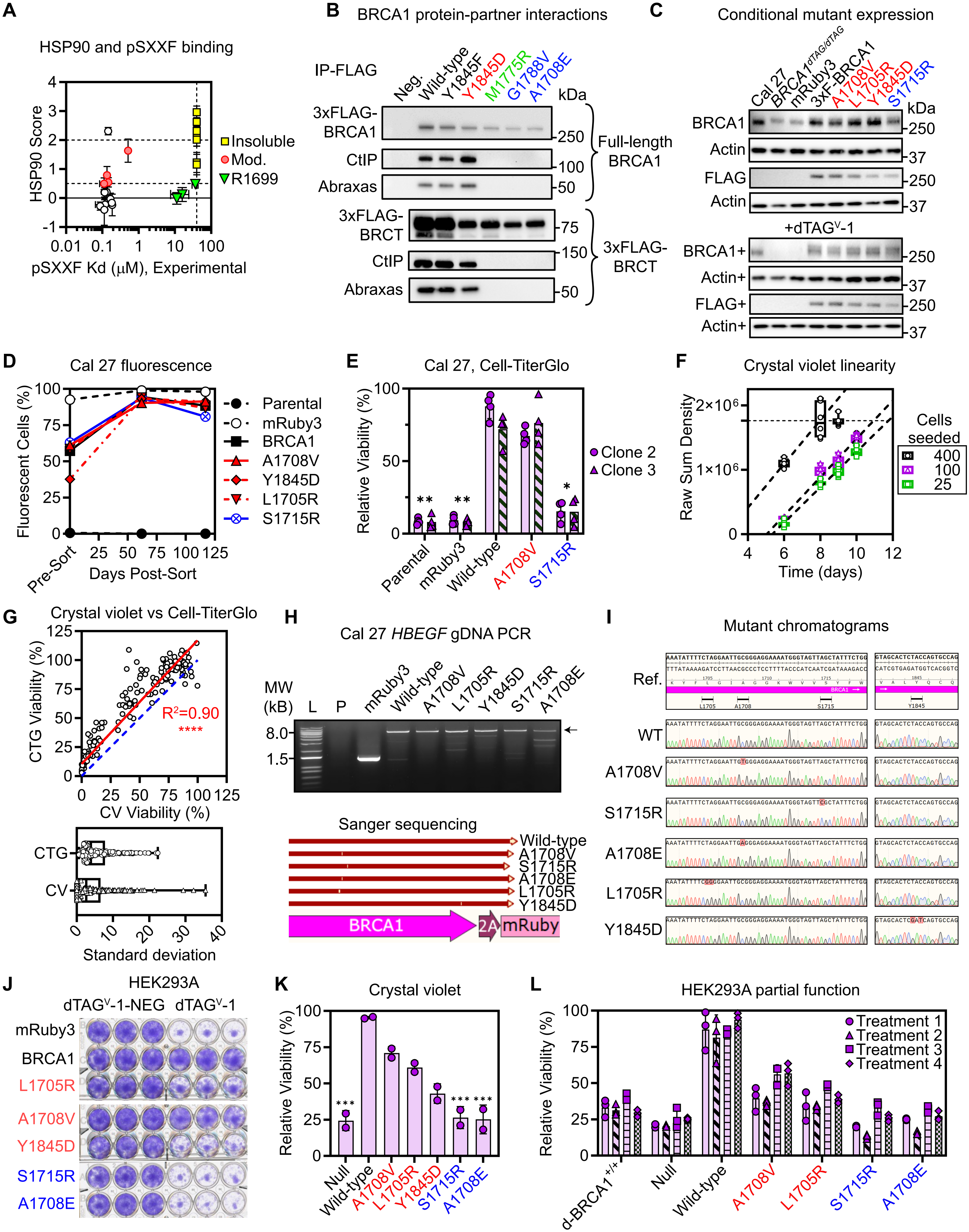


**Figure S2. Moderate HSP90 engagement signifies maintenance of BRCA1 function, Related to Figure 2**

### Figure S3

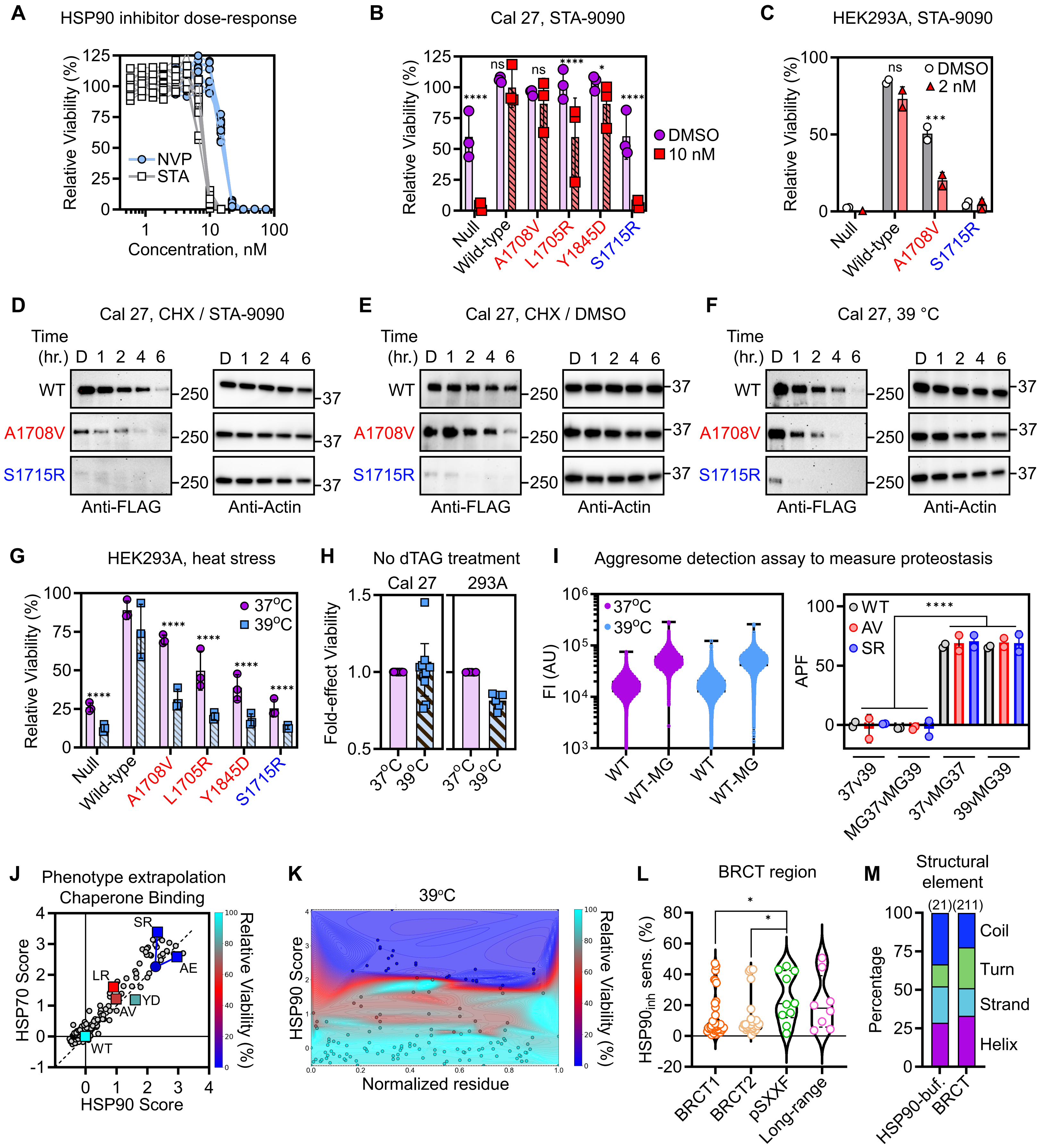


**Figure S3. HSP90 stabilizes BRCA1 variants in cancer cells, Related to Figure 3**
