## Supplementary material for "HSP90 buffers deleterious genetic variations in *BRCA1*": Figure S4

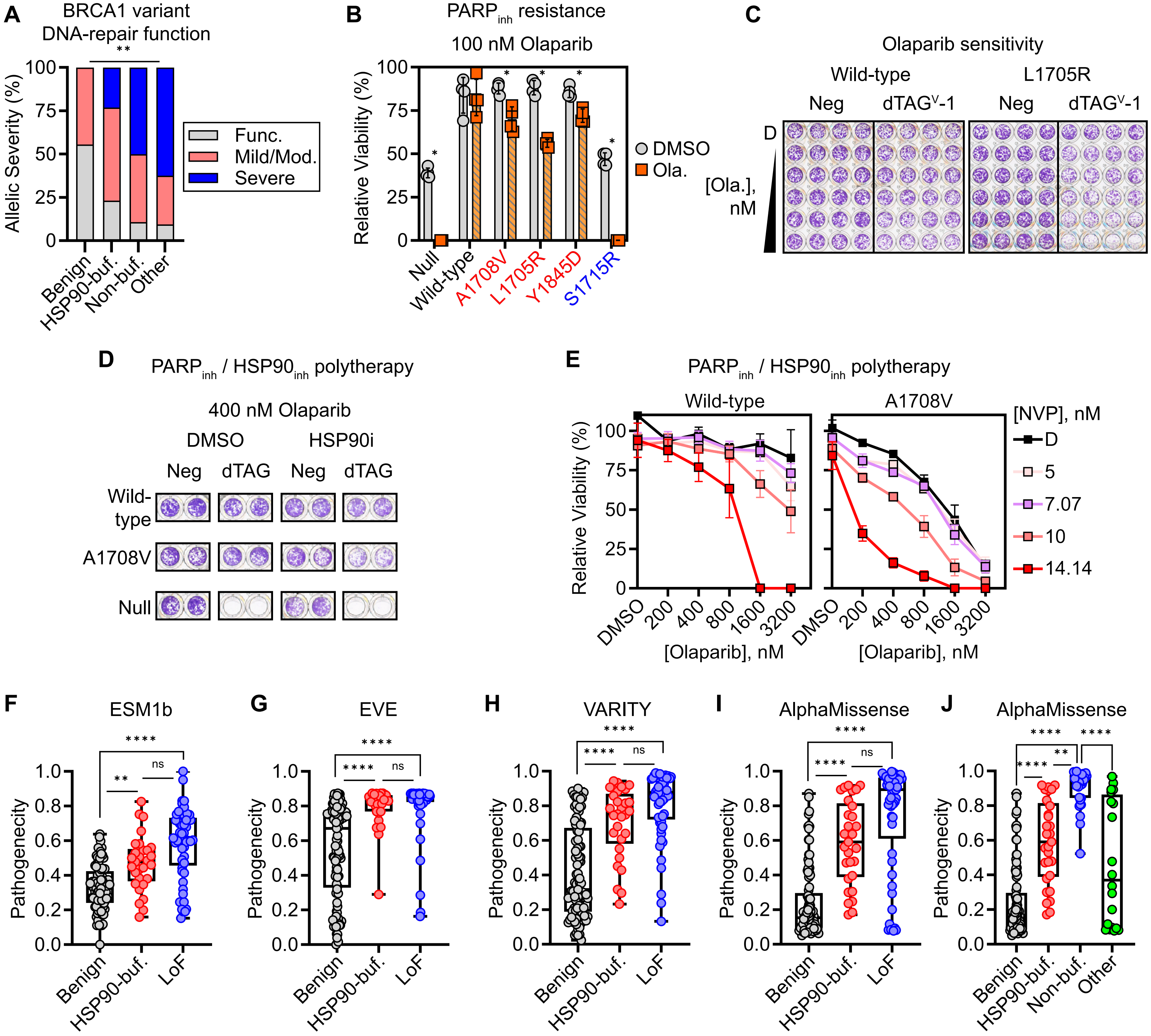


**Figure S4. HSP90-buffered BRCA1 variants exhibit distinct pathogenic properties, Related to Figure 4**
